# Short-term monocular deprivation engages rapid, inhibition-gated ocular dominance plasticity in mouse visual cortex

**DOI:** 10.64898/2026.04.13.718078

**Authors:** Irene di Marco, Gabriele Sansevero, Nicoletta Berardi, Alessandro Sale

## Abstract

Ocular dominance (OD) plasticity has long served as a canonical model of experience-dependent cortical plasticity, traditionally thought to be confined to a developmental critical period. Recent human studies, however, show that brief monocular deprivation of just a few hours induces rapid and fully reversible OD shifts, revealing a form of homeostatic plasticity that persists into adulthood. The underlying cellular and circuit mechanisms remain unknown, largely due to the lack of suitable preclinical models. Here, we establish and validate a mouse model of short-term monocular deprivation that closely recapitulates the temporal dynamics observed in humans. Using in vivo electrophysiology in awake mice, we show that two hours of monocular deprivation induce robust yet reversible OD shifts in adult visual cortex, and even larger shifts during the critical period. These shifts are driven by reciprocal modulation of eye-specific cortical responses, with enhanced visual evoked potentials from the deprived eye and concurrent suppression of the non-deprived eye. Chemogenetic manipulation of parvalbumin-positive (PV) interneurons reveals a causal, bidirectional role for PV-mediated inhibition in gating this plasticity: transient PV suppression amplifies OD shifts to juvenile-like levels, whereas PV enhancement constrains or abolishes them. Together, these findings identify a fast, inhibition-gated form of homeostatic OD plasticity operating across developmental stages. This tractable model bridges human perceptual plasticity with defined circuit mechanisms and offers a foundation for developing translational strategies for visual disorders such as amblyopia.

## Introduction

The primary visual cortex (V1) exhibits a striking form of experience-dependent plasticity early in postnatal life, known as ocular dominance (OD) plasticity, which has long served as a canonical model for studying cortical development.

This phenomenon was first described by Hubel and Wiesel, who showed that monocular deprivation (MD) during a defined critical period (CP) shifts cortical responsiveness toward the non-deprived eye (Hubel & Wiesel, 1959, 1970). These studies established V1 as a central model for understanding how sensory experience shapes synaptic organization. Since then, MD-induced OD plasticity has become a key paradigm for studying the cellular and molecular bases of cortical plasticity. This work has clarified the roles of inhibitory circuits, structural remodeling, and the timing of developmental CPs (Espinosa & Stryker, 2012; Hensch, 2005; Levelt & Lubener, 2012; Rittenhouse et al., 1999).

OD plasticity was traditionally considered largely confined to the juvenile CP. Recent studies in adult humans have shown that short-term monocular deprivation (STMD) causes a counterintuitive shift in OD, enhancing the contribution of the deprived eye (Lunghi et al., 2011, 2013). This effect reveals a rapid form of homeostatic plasticity in the mature visual cortex. About 150 minutes of visual deprivation, achieved by covering one eye with a translucent patch, increases the perceptual dominance of the deprived eye. This effect has been reliably measured using binocular rivalry in both adults (Binda & Lunghi, 2017; Lunghi, Berchicci, et al., 2015; Lunghi, Emir, et al., 2015; Lunghi et al., 2011, 2013; Lunghi & Sale, 2015; Menicucci et al., 2022; Prosper et al., 2023) and adolescents (Nguyen et al., 2023). The effect is transient, usually lasting 90–120 minutes after patch removal (Lunghi et al., 2013), but non-REM sleep can extend its duration for several hours (Menicucci et al., 2022).

This homeostatic shift is not limited to perception. Pattern-onset visual evoked potentials (VEPs) reveal an enhanced C1 amplitude for the deprived eye and reduced responses for the non-deprived eye, together with increased alpha-band activity (Lunghi, Berchicci, et al., 2015). Magnetic resonance spectroscopy further shows that STMD decreases resting GABA levels in V1, indicating a transient disinhibitory state that allows rapid plasticity (Lunghi, Emir, et al., 2015).

Despite the evidence in humans, the neural and circuit mechanisms underlying STMD remain unclear, largely because an appropriate animal model is lacking. In this study, we establish and validate a mouse model of STMD that faithfully replicates the human experimental protocol. Wild-type mice were subjected to 2 hours of MD either during adulthood or at the CP (postnatal day 28, P28), In vivo VEP recordings were performed in awake animals immediately before and after deprivation. The transience of the effect was assessed with a third VEP recording, performed 2 hours after reopening of the deprived eye. Lastly, to investigate the contribution of inhibitory circuitry, we selectively modulated the activity of parvalbumin-positive (PV) interneurons using chemogenetic tools.

This model offers a robust platform for dissecting the cellular and circuit mechanisms underlying rapid homeostatic plasticity and provides new opportunities for translational research on visual disorders such as amblyopia.

## Methods

### Animal treatment

C57BL/6J and B6;129P2-Pvalbtm1(cre)Arbr/J mice were used in accordance with the guidelines of the Italian Ministry of Public Health. All procedures were approved by the Ministry (protocols 23C and 24C) and were designed to minimize animal use and discomfort. Mice were housed at 21 °C under a 12-h light/dark cycle with ad libitum access to food and water.

### Headplate implantation

Adult wild-type C57BL/6J (∼P60), B6;129P2-Pvalbtm1(cre)Arbr/J (∼P60), and young wild-type C57BL/6J (P21) mice were anesthetized via intraperitoneal injection of Zolazepam + Tiletamine (Zoletil-100, 40 mg/kg, Virbac) and Xylazine (10 mg/kg, Sigma) and secured in a stereotaxic apparatus. A cranial window was made over the binocular zone of V1 (1.5–3.5 mm lateral to lambda in adults; 1.7–2.2 mm in P21 mice). A metal headplate was fixed to the parietal bone contralateral to the window using dental resin cement (Super-Bond, C&B). The cranial window was filled with agar to maintain cortical moisture and covered with silicone elastomer (Kwik-Cast, World Precision Instruments) to protect the tissue. A screw in the cerebellum served as a ground reference, and dental cement (Paladur®) secured the exposed bone. Body temperature was maintained with a feedback-regulated heating pad, and mice were closely monitored post-surgery. After one week of recovery, animals were habituated to the recording apparatus.

### In vivo electrophysiology

To investigate the effects of short-term monocular deprivation (STMD) and its underlying mechanisms, we adapted a human protocol (Lunghi et al., 2011, 2013) and compared deprivation of the dominant eye (contralateral to the recorded hemisphere, CONTRA) versus the non-dominant eye (ipsilateral, IPSI). Awake, head-fixed mice (∼P74 or P28) underwent acute electrophysiological recordings from the binocular zone of V1 at three time points: pre-deprivation (Pre), immediately after 2 hours of monocular deprivation (Post), and 2 hours after eye reopening (After 2h).

Local field potentials were recorded using a 16-channel silicon probe (A1x16-3 mm-50-703-A16, Neuronexus Technologies) connected to a 16-channel Open Ephys acquisition system. Signals were filtered between 0.3 and 275 Hz and sampled at 30.3 kHz. Data analysis was performed using custom Python scripts. Visual stimuli were generated in Python using the PsychoPy extension and presented on a monitor (Asus VG278QR, 165 Hz refresh rate, 400 cd m−2 mean luminance) positioned 20 cm from the animal. VEPs were evoked using sinusoidal gratings (0.06 cycles/deg) with abrupt phase reversals (2 Hz temporal frequency) and analyzed in the time domain to measure peak-to-baseline amplitude and latency.

Ocular dominance was quantified using the contralateral/ipsilateral (C/I) VEP ratio, calculated as the ratio of VEP amplitudes evoked through the contralateral versus ipsilateral eye. Manual shutters were used to alternately close each eye during recordings.

### Short-term monocular deprivation

Mice were anesthetized with isoflurane (4% for induction, 1.5–3.5% for maintenance). Monocular deprivation (MD) or binocular deprivation (BD, in control animals) was performed by suturing the eyelid with 7-0 silk. During the 2 hours of MD, we continuously monitored the animals to assess the absence of eyelid reopening and the maintenance of wakefulness. After 2 hours, the eyelids were reopened under isoflurane anesthesia (4% induction, 1.5–3.5% maintenance). In control experiments, the animals underwent the same isoflurane anesthesia procedure (4% induction, 1.5–3.5% maintenance) without MD. Animals were allowed to fully recover from anesthesia before undergoing electrophysiological recordings.

### Intracortical AAV injection in V1

Adult PV-Cre mice (∼P60; B6;129P2-Pvalbtm1(cre)Arbr/J, Jackson Laboratories) or adult C57BL/6J mice were anesthetized with Zolazepam + Tiletamine (Zoletil-100, 40 mg/kg, Virbac) and Xylazine (10 mg/kg, Sigma) and mounted on a stereotaxic apparatus. The skin above the skull was removed, and a small hole was drilled over V1 (2.8 mm lateral to lambda).

For selective suppression/enhancement of PV interneuron activity, PV-Cre mice received bilateral injections of AAV8-hSyn-DIO-hM4D(Gi)-mCherry/AAV8-hSyn-DIO-hM3D(Gq)-mCherry (Addgene). To suppress V1 activity more broadly, C57BL/6J mice were injected monolaterally with AAV8-hSyn-HA-hM4D(Gi)-mCherry (Addgene #50475-AAV8) in the hemisphere contralateral to the recording site (see Supplementary Figs. 1–2).

Viral injections were delivered using a 10 μL Hamilton syringe connected to a nanoliter syringe pump (Kd Scientific) at a speed of 0.05 μL/min. Injections were made at two cortical depths (220 μm and 450 μm below the pial surface) at the same insertion site (200 nL per depth). Each injection lasted 4 minutes per depth, starting from the deeper site, followed by a 2-minute wait before withdrawing the syringe.

Following surgery, body temperature was maintained with a feedback-regulated heating pad, and mice were carefully monitored during recovery. Headplate implantation for electrophysiological recordings was performed as described above. Habituation to the apparatus began one week after surgery, and experimental protocols were initiated two weeks after viral injection.

### Chemogenetic experiments

Clozapine N-oxide (CNO; Tocris Bioscience, Cat. No. 4936) was dissolved in sterile 0.9% NaCl. For chemogenetic manipulation, two intraperitoneal injections of CNO (2 mg/kg) were administered. In PV-Cre mice, CNO was used to suppress PV interneuron activity in V1, whereas in C57BL/6J mice it was used to suppress V1 activity more broadly during the 2-hour monocular deprivation. The first injection was given immediately after eyelid closure, and the second at the end of the first hour of deprivation.

### Immunohistochemistry

To verify AAV expression in PV interneurons, PV-Cre mice were perfused transcardially with PBS followed by 4% paraformaldehyde (PFA) in phosphate buffer. Brains were post-fixed for 24 hours at 4°C, then immersed in 30% sucrose in phosphate buffer. Coronal sections (50 µm) were cut on a freezing microtome and collected in PBS. Free-floating sections were blocked in 10% bovine serum with 0.2% Triton X-100 for 1.5 hours at room temperature, followed by overnight incubation at 4°C with guinea pig anti-parvalbumin primary antibody (1:1000; SySy, Germany) diluted in blocking solution. Sections were washed three times with PBS and incubated for 2 hours at room temperature with Alexa488-conjugated anti-rabbit secondary antibody (1:500; Invitrogen). Nuclei were counterstained with Hoechst (1:500; Sigma-Aldrich) for 15 minutes at room temperature. Sections were mounted on glass slides and covered with VectaShield mounting medium, and fluorescence was acquired using a confocal microscope.

For assessing AAV expression in both PV-Cre and C57BL/6J mice, brains were post-fixed overnight at 4°C and immersed in 30% sucrose in phosphate buffer. Coronal sections (50 µm) were cut, washed in PBS-T (0.3% Triton), incubated with Hoechst (1:500) for 15 minutes at room temperature, mounted on slides, and coverslipped with VectaShield. Fluorescence images were acquired on a fluorescence microscope equipped with an Apotome 2.0 slit.

### Statistical analysis

All statistical analyses were performed using R (version 4.1.2). Data were first tested for normality. For normally distributed data, parametric tests were applied; if normality was not met, non-parametric tests were used. Differences between two independent groups were assessed with a two-tailed unpaired t-test, while comparisons between two dependent groups were performed with a two-tailed paired t-test. For multiple group comparisons, one-way ANOVA, repeated measures (RM) one-way ANOVA, two-way ANOVA, or two-way RM ANOVA were applied as appropriate. Non-normally distributed data from more than two groups were analyzed using one-way ANOVA on ranks or two-way ANOVA on ranks. Data are expressed as mean ± SEM, and sample sizes (n) are reported for each experiment.

## Results

### 1. Short-term monocular deprivation induced transient OD shifts in adult mice

To investigate the effects of STMD and its underlying mechanisms, we adapted the human protocol (Lunghi et al., 2011, 2013), comparing deprivation of the dominant eye (contralateral to the recorded hemisphere, CONTRA) versus the non-dominant eye (ipsilateral, IPSI). Awake, head-fixed mice (∼postnatal day 74, P74) underwent acute electrophysiological recordings from the binocular zone of V1 at three time points: pre-deprivation (Pre), immediately after 2 hours of CONTRA or IPSI eye deprivation (Post), and 2 hours after eye reopening (After 2h; **Figure 1A**,**B**).

**Figure 1.**
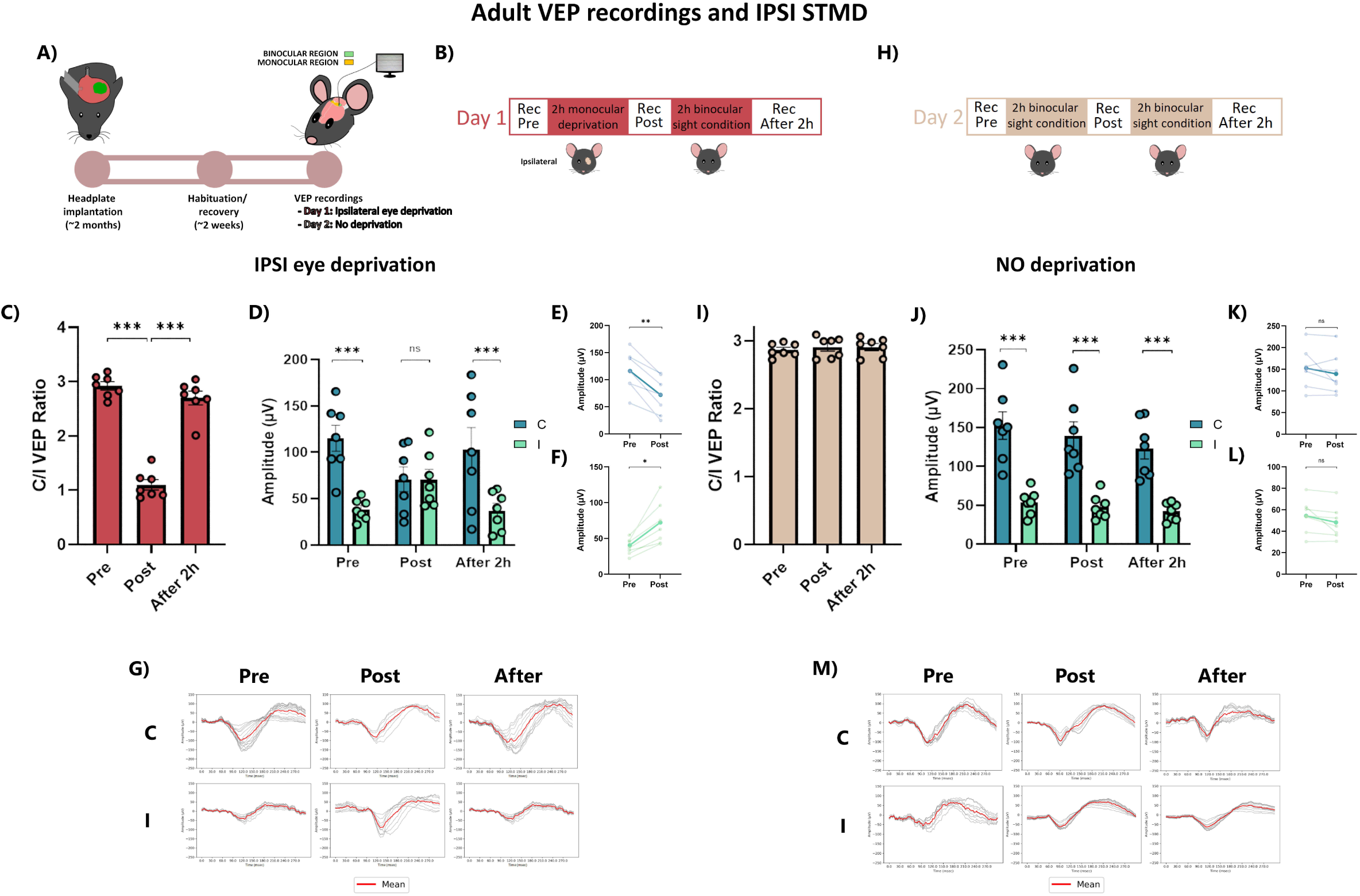
IPSI short-term monocular deprivation (STMD) induces rapid and reversible ocular dominance plasticity in adult mice. **a**, Schematic of the experimental protocol for IPSI STMD. **b,** Timeline of electrophysiological recordings on Day 1: baseline (Pre), immediately after STMD (Post), and after 2 h of restored binocular vision (After 2h). **c,** Contralateral/ipsilateral (C/I) VEP ratio in response to a low spatial frequency grating (0.06 cycles/degree). The C/I ratio significantly decreased immediately after IPSI STMD compared to baseline (one-way repeated-measures ANOVA, p < 0.001) and returned to baseline after 2 h of binocular vision (p = 0.209). **d,** Mean VEP amplitudes for contralateral and ipsilateral eye stimulation across all time points. **e,** Mean contralateral-eye VEP amplitudes before and after IPSI STMD. Monocular deprivation induced a significant reduction in contralateral responses (two-way repeated-measures ANOVA, p < 0.01). **f,** Mean ipsilateral-eye VEP amplitudes before and after IPSI STMD. Monocular deprivation induced a significant increase in ipsilateral responses (two-way repeated-measures ANOVA, p < 0.01). In **e,f**, darker lines indicate the mean across animals. **g,** Representative VEP traces from a single animal across time points. **h,** Timeline of electrophysiological recordings on Day 2 (control condition without deprivation). **i,** Three consecutive recordings without monocular deprivation did not induce significant changes in the C/I VEP ratio (one-way repeated-measures ANOVA, p = 0.489). **j,** Mean VEP amplitudes for contralateral and ipsilateral eye stimulation across control time points. **k,** Mean contralateral-eye VEP amplitudes at baseline (Pre) and after 2 h of binocular vision (Post) in control recordings (two-way repeated-measures ANOVA, p = 0.845). **l,** Mean ipsilateral-eye VEP amplitudes at baseline and after 2 h of binocular vision in control recordings (two-way repeated-measures ANOVA, p = 0.994). In **k,l**, darker lines indicate the mean. **m,** Representative VEP traces from a single animal across control time points. C, contralateral eye; I, ipsilateral eye; Pre, before deprivation; Post, immediately after deprivation; After, 2 h after deprivation. Error bars indicate s.e.m. Statistical significance: ns, not significant; *p < 0.05; **p < 0.01; ***p < 0.001.

OD was assessed by calculating the contralateral-to-ipsilateral (C/I) VEP amplitude ratio. Under baseline conditions, the C/I VEP ratio is approximately 2.7–3, (Kaplan et al., 2016; Mazziotti et al., 2017; Porciatti et al., 1999).

#### 1.1. IPSI eye deprivation in adult mice

The baseline C/I VEP ratio in adult mice (n = 7; ∼P74) was 2.923 ± 0.075. IPSI STMD caused a significant reduction in the C/I VEP ratio (One-Way RM ANOVA, DF = 2, F = 129.8; Tukey method, Pre = 2.923 ± 0.075 vs. Post = 1.101 ± 0.093, p < 0.001), which returned to baseline after 2 hours of binocular viewing (One-Way RM ANOVA, Pre = 2.923 ± 0.075 vs. After 2h = 2.7 ± 0.124, p = 0.209; **Figure 1C**), indicating the transient nature of this plasticity.

To dissect the contributions of each eye, individual VEP amplitudes were analyzed. STMD induced a significant decrease in CONTRA VEP amplitudes (Two-Way RM ANOVA, eye × time, DF = 2, F = 13.08; Tukey method, Pre = 115.157 ± 13.94 μV vs. Post = 70.977 ± 13.28 μV, p < 0.01; **Figure 1E,G**), accompanied by a significant increase in IPSI VEP amplitudes (Pre = 38.377 ± 4.25 μV vs. Post = 70.292 ± 11.208 μV, p < 0.05; **Figure 1F,G**). Before deprivation, CONTRA responses were significantly stronger than IPSI (Contra Pre = 115.157 ± 13.94 μV vs. Ipsi Pre = 38.377 ± 4.25 μV, p < 0.001), consistent with typical OD. This asymmetry was abolished immediately after STMD (Contra Post = 70.977 ± 13.28 μV vs. Ipsi Post = 70.292 ± 11.208 μV, p = 1.00). Following 2 hours of binocular vision, VEP amplitudes returned to their original asymmetry (Contra After 2h = 102.901 ± 23.71 μV vs. Ipsi After 2h = 37.088 ± 7.78 μV, p < 0.001; **Figure 1D,G**).

As a control, the same animals (n = 7) were recorded on the following day at three consecutive time-points (Pre, Post, After 2h; **Figure 1A,H**) without any eye deprivation, confirming that changes in the C/I VEP ratio were specifically induced by IPSI STMD rather than repeated recordings. No significant changes in the C/I VEP ratio were observed (Pre = 2.868 ± 0.036, Post = 2.909 ± 0.057, After 2h = 2.910 ± 0.051; One-way ANOVA, DF = 2, F = 0.761, Tukey method, p = 0.489; **Figure 1I**). Repeated recordings similarly did not significantly alter CONTRA (Contra Pre = 152.68 ± 17.67 μV vs. Contra Post = 139.43 ± 17.99 μV; Two-Way RM ANOVA, eye × time, DF=2, F=0.566, Tukey method, p = 0.845; **Figure 1K,M**) or IPSI VEP amplitudes (Ipsi Pre = 54.09 ± 5.993 μV vs. Ipsi Post = 48.038 ± 5.822 μV; Two-Way RM ANOVA, p = 0.994; **Figure 1L,M**). The C/I amplitude difference remained significant across all three recordings (Two-Way RM ANOVA, p < 0.001; **Figure 1H,M**). These findings confirm that the experimental protocol, including head-fixation and repeated recordings, did not affect OD or VEP amplitudes.

#### 1.2 CONTRA eye deprivation in adult mice

To further examine the effects of STMD on visual cortical responses, we conducted a complementary experiment in a separate group of awake, head-fixed mice (n = 9; ∼P74), using the same three-session protocol (Pre, Post, After 2h; **Figure 2A,B**). Here, the eye contralateral to the recording site was deprived.

**Figure 2.**
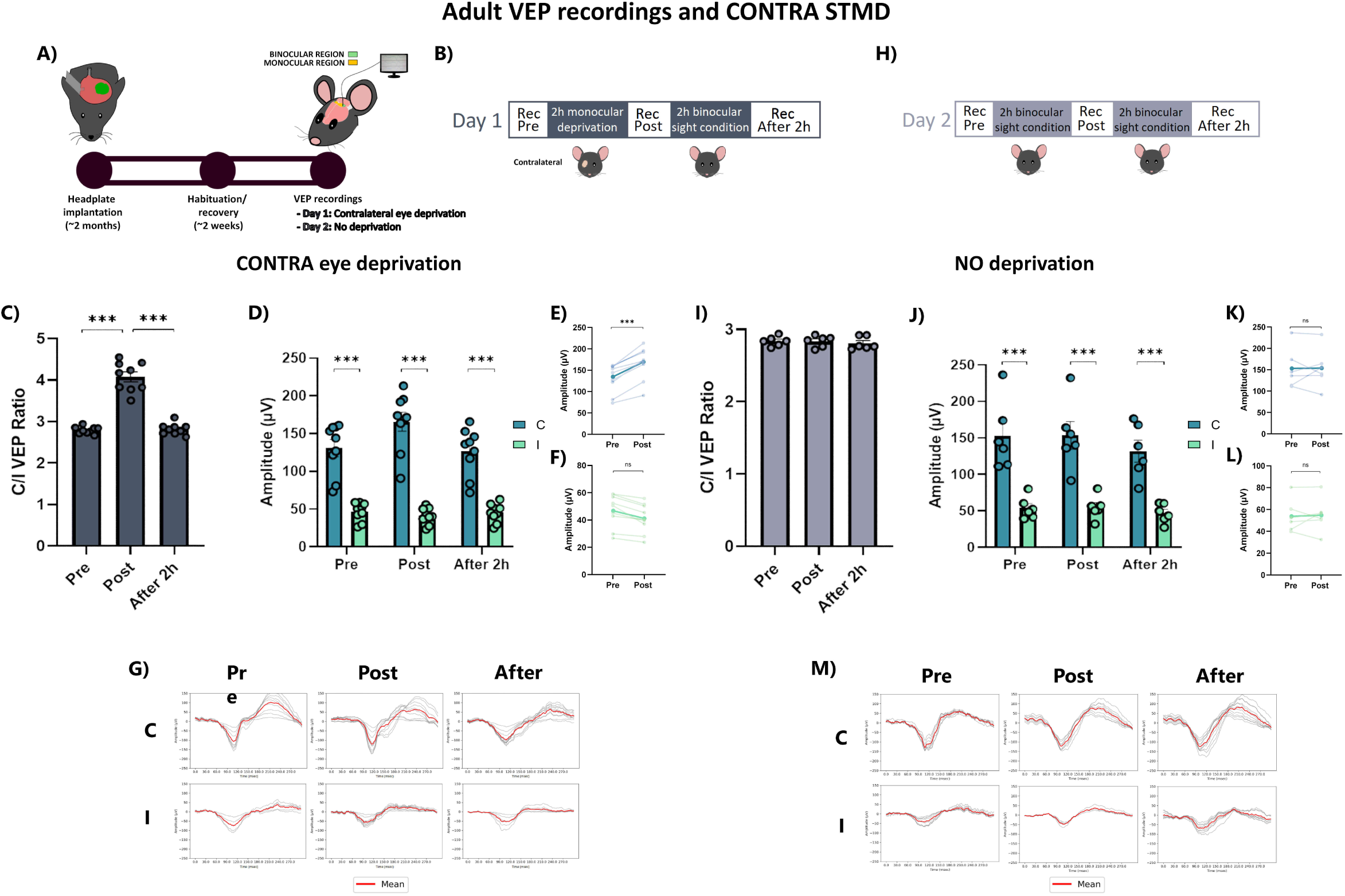
CONTRA short-term monocular deprivation (STMD) induces rapid and reversible ocular dominance plasticity in adult mice. **a**, Schematic of the experimental protocol for CONTRA STMD. **b,** Timeline of electrophysiological recordings on Day 1: baseline (Pre), immediately after STMD (Post), and after 2 h of restored binocular vision (After 2h). **c,** Contralateral/ipsilateral (C/I) VEP ratio in response to a low spatial frequency grating (0.06 cycles/degree). The C/I ratio significantly increased immediately after CONTRA STMD compared to baseline (one-way repeated-measures ANOVA, p < 0.001) and returned to baseline after 2 h of binocular vision (p = 0.962). **d,** Mean VEP amplitudes for contralateral and ipsilateral eye stimulation across all time points. **e,** Mean contralateral-eye VEP amplitudes before and after CONTRA STMD. Monocular deprivation induced a significant increase in contralateral responses (two-way repeated-measures ANOVA, p < 0.001). **f,** Mean ipsilateral-eye VEP amplitudes before and after CONTRA STMD. Monocular deprivation did not significantly affect ipsilateral responses (two-way repeated-measures ANOVA, p = 0.936). In **e,f**, darker lines indicate the mean across animals. **g,** Representative VEP traces from a single animal across time points. **h,** Timeline of electrophysiological recordings on Day 2 (control condition without deprivation). **i,** Three consecutive recordings without monocular deprivation did not induce significant changes in the C/I VEP ratio (one-way repeated-measures ANOVA, p = 0.621). **j,** Mean VEP amplitudes for contralateral and ipsilateral eye stimulation across control time points. **k,** Mean contralateral-eye VEP amplitudes at baseline (Pre) and after 2 h of binocular vision (Post) in control recordings (two-way repeated-measures ANOVA, p = 0.999). **l,** Mean ipsilateral-eye VEP amplitudes at baseline and after 2 h of binocular vision in control recordings (two-way repeated-measures ANOVA, p = 0.999). In **k,l**, darker lines indicate the mean. **m,** Representative VEP traces from a single animal across control time points. C, contralateral eye; I, ipsilateral eye; Pre, before deprivation; Post, immediately after deprivation; After, 2 h after deprivation. Error bars indicate s.e.m. Statistical significance: ns, not significant; *p < 0.05; **p < 0.01; ***p < 0.001.

The baseline C/I VEP ratio was 2.796 ± 0.027. CONTRA STMD produced a significant increase (One-Way RM ANOVA, DF = 2, F = 117.6; Tukey method, Pre = 2.796 ± 0.027 vs. Post = 4.080 ± 0.117, p < 0.001). This effect was transient, returning to baseline after 2 hours of binocular vision (One-Way RM ANOVA, Pre = 2.796 ± 0.027 vs. After 2h = 2.821 ± 0.044, p = 0.962; **Figure 2C**).

Examining individual VEP amplitudes, CONTRA responses increased significantly after deprivation (Pre = 131.126 ± 11.18 μV vs. Post = 165.936 ± 12.57 μV; Two-Way RM ANOVA, eye × time, DF = 2, F = 9.748; Tukey method, p < 0.001; **Figure 2E,G**), while IPSI VEP amplitudes showed a concurrent, non-significant decrease (Pre = 46.912 ± 3.99 μV vs. Post = 41.315 ± 3.58 μV, p = 0.936; **Figure 2F,G**). The C/I amplitude difference remained significant across all sessions (Two-Way RM ANOVA, p < 0.001; **Figure 2D,G**).

As a control, a subset of animals (n = 6) underwent the same three-session recording protocol without visual deprivation (Pre, Post, After 2h; **Figure 2A,H**). No significant changes in the C/I VEP ratio were detected (Pre = 2.838 ± 0.027, Post = 2.837 ± 0.031, After 2h = 2.809 ± 0.035, One-Way RM ANOVA, DF = 2, F = 0.5; Tukey method, p = 0.621; **Figure 2I**). Repeated recordings also did not significantly alter CONTRA (Pre = 152.74 ± 17.77 μV vs. Post = 153.57 ± 17.32 μV; Two-way ANOVA RM, eye × time, DF=2, F=190.121, Tukey method, p = 0.999; **Figure 2K,M**) or IPSI responses (Pre = 53.96 ± 5.64 μV vs. Post = 54.93 ± 5.88 μV; Two-way ANOVA RM, p = 0.999; **Figure 2L,M**). Contralateral eye dominance remained significant across all sessions (Two-Way RM ANOVA, p < 0.001; **Figure 2H,M**).

Overall, these results show that STMD in adult mice produces a rapid but transient OD shift in V1, mediated by reciprocal modulation of eye-specific cortical responses.

### 2. Juvenile visual cortex exhibits stronger transient OD plasticity after short-term monocular deprivation

To investigate whether STMD triggers similar visual cortical plasticity during development, we conducted parallel experiments in juvenile mice at postnatal day 28 (P28), the peak of the critical period CP. Two separate groups underwent either IPSI or CONTRA eye deprivation. The experimental protocol mirrored that used in adult mice, with awake, head-fixed animals undergoing acute VEP recordings from the binocular zone of V1 at three time points: pre-deprivation (Pre), immediately after 2 hours of CONTRA or IPSI eye deprivation (Post), and 2 hours after eye reopening (After 2h; **Figure 3A,B**; **Figure 4A,B**).

**Figure 3.**
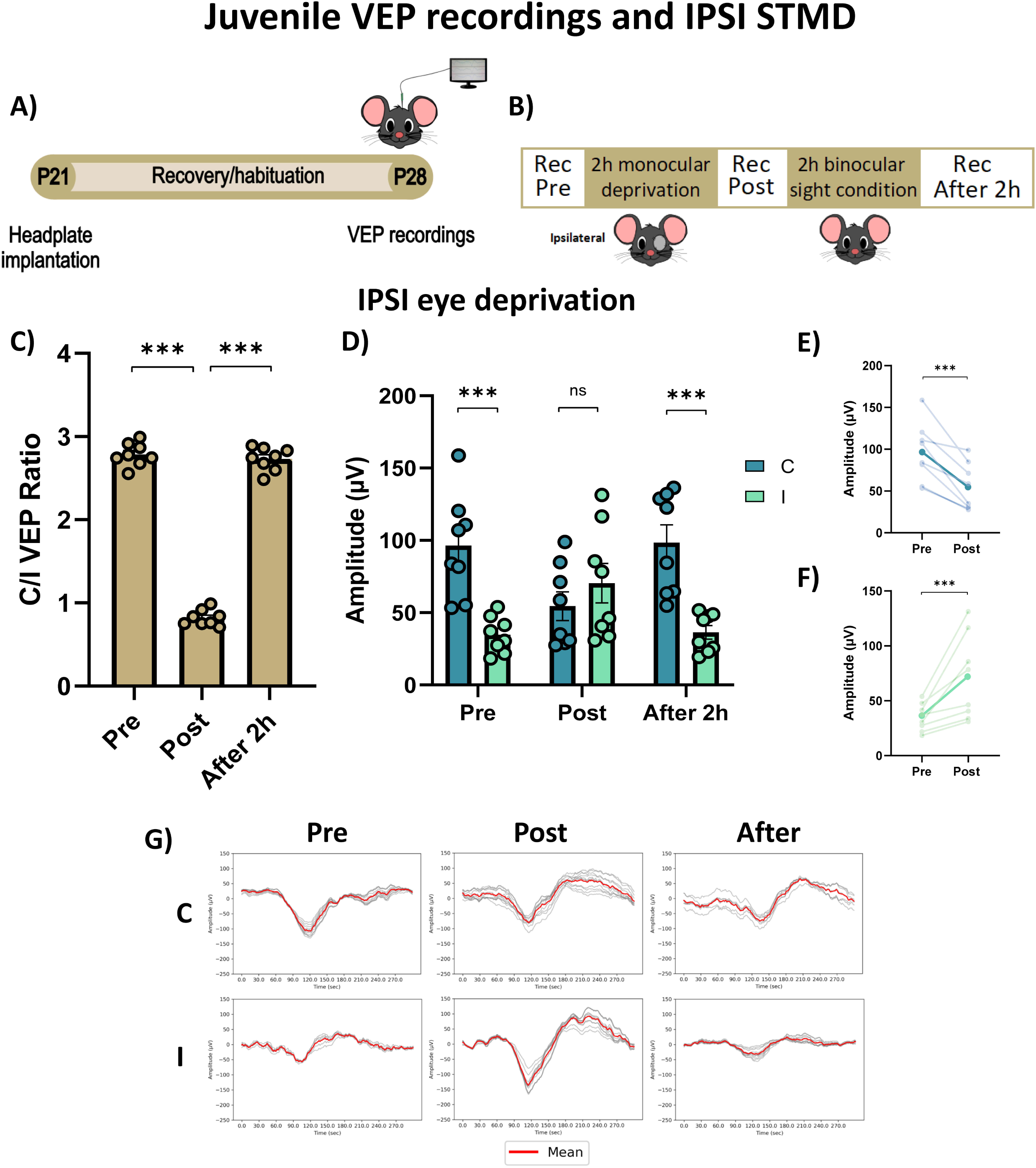
IPSI short-term monocular deprivation (STMD) induces enhanced ocular dominance plasticity in juvenile mice. **a**, Schematic of the experimental protocol for IPSI STMD. **b,** Timeline of electrophysiological recordings: baseline (Pre), immediately after STMD (Post), and after 2 h of restored binocular vision (After 2h). **c,** Contralateral/ipsilateral (C/I) VEP ratio in response to a low spatial frequency grating (0.06 cycles/degree). The C/I ratio significantly decreased immediately after IPSI STMD compared to baseline (one-way repeated-measures ANOVA, p < 0.001) and returned to baseline after 2 h of binocular vision (p = 0.604). **d,** Mean VEP amplitudes for contralateral and ipsilateral eye stimulation across all time points. **e,** Mean contralateral-eye VEP amplitudes before and after IPSI STMD. Monocular deprivation induced a significant reduction in contralateral responses (two-way repeated-measures ANOVA, p < 0.001). **f,** Mean ipsilateral-eye VEP amplitudes before and after IPSI STMD. Monocular deprivation induced a significant increase in ipsilateral responses (two-way repeated-measures ANOVA, p < 0.001). In **e,f**, darker lines indicate the mean across animals. **g,** Representative VEP traces from a single animal across time points. C, contralateral eye; I, ipsilateral eye; Pre, before deprivation; Post, immediately after deprivation; After, 2 h after deprivation. Error bars indicate s.e.m. Statistical significance: ns, not significant; *p < 0.05; **p < 0.01; ***p < 0.001.

**Figure 4.**
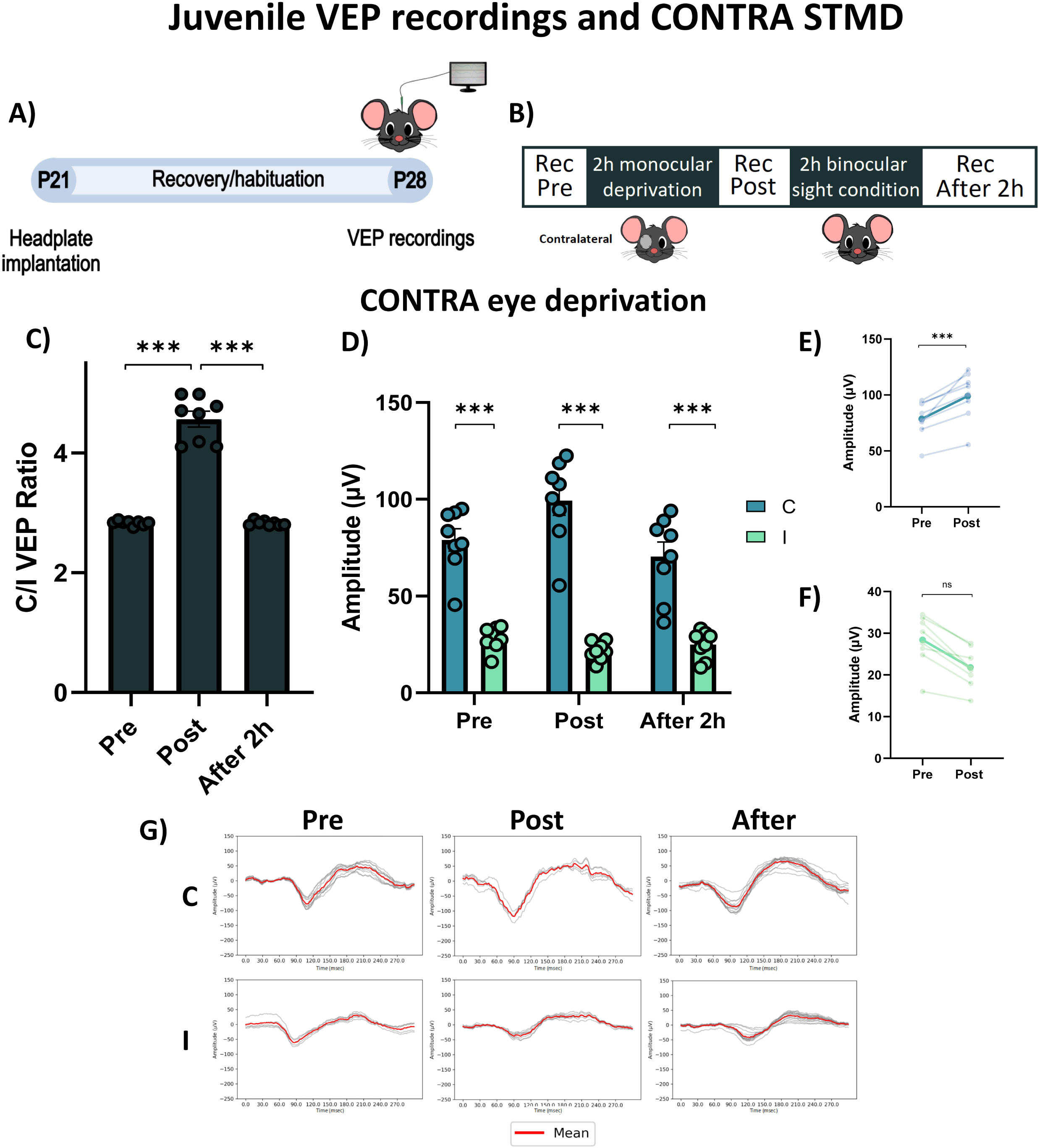
CONTRA short-term monocular deprivation (STMD) induces ocular dominance plasticity in juvenile mice. **a**, Schematic of the experimental protocol for CONTRA STMD. **b,** Timeline of electrophysiological recordings: baseline (Pre), immediately after STMD (Post), and after 2 h of restored binocular vision (After 2h). **c,** Contralateral/ipsilateral (C/I) VEP ratio in response to a low spatial frequency grating (0.06 cycles/degree). The C/I ratio significantly increased immediately after CONTRA STMD compared to baseline (one-way repeated-measures ANOVA, p < 0.001) and returned to baseline after 2 h of binocular vision (p = 0.998). **d,** Mean VEP amplitudes for contralateral and ipsilateral eye stimulation across all time points. **e,** Mean contralateral-eye VEP amplitudes before and after CONTRA STMD. Monocular deprivation induced a significant increase in contralateral responses (two-way repeated-measures ANOVA, p < 0.001). **f,** Mean ipsilateral-eye VEP amplitudes before and after CONTRA STMD. Monocular deprivation induced a non-significant decrease in ipsilateral responses (two-way repeated-measures ANOVA, p = 0.689). In **e,f**, darker lines indicate the mean across animals. **g,** Representative VEP traces from a single animal across time points. C, contralateral eye; I, ipsilateral eye; Pre, before deprivation; Post, immediately after deprivation; After, 2 h after deprivation. Error bars indicate s.e.m. Statistical significance: ns, not significant; *p < 0.05; **p < 0.01; ***p < 0.001.

#### 2.1 IPSI eye deprivation in juvenile mice

In juvenile mice (n = 8; P28), the baseline C/I VEP ratio was 2.783 ± 0.049. IPSI STMD induced a marked reduction in the ratio (Pre = 2.783 ± 0.049 vs. Post = 0.819 ± 0.033, One-Way RM ANOVA, DF = 2, F = 879.1; Tukey method, p < 0.001). This effect was transient, returning to baseline after 2 hours of binocular vision (Pre = 2.783 ± 0.049 vs. After 2h = 2.731 ± 0.049, One-Way RM ANOVA, p = 0.604; **Figure 3C**).

The IPSI-induced OD shift was significantly larger in juveniles than in adults, reflected by a lower post-deprivation C/I VEP ratio in P28 mice (two-tailed unpaired t-test, Post adult = 1.101 ± 0.093 vs. Post P28 = 0.819 ± 0.033; t = 2.843, DF = 7.57, p < 0.05; **Figure 5A**).

**figure 5.**
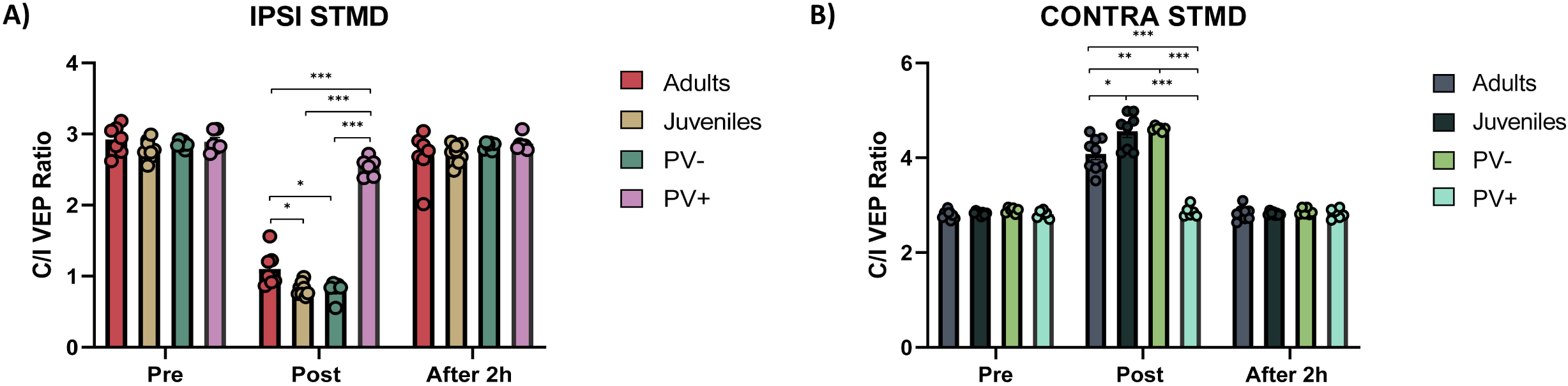
Comparative analysis of IPSI and CONTRA short-term monocular deprivation (STMD) across experimental groups. **a**, IPSI STMD induced a significantly greater effect in juveniles and PV- mice compared to adult animals (two-tailed unpaired t-test, Post adults vs. Post juveniles, p < 0.05; Post adults vs. Post PV-, p < 0.01). In contrast, PV+ mice exhibited a significantly reduced IPSI STMD effect relative to all other groups (Post adults vs. PV+, p < 0.001; Post PV-vs. PV+, p < 0.001; Post juveniles vs. PV+, p < 0.001). **b,** CONTRA STMD produced a stronger effect in juveniles and PV- mice compared to adult animals (Post adults vs. Post juveniles, p < 0.05; Post adults vs. Post PV-, p < 0.05). PV+ mice showed a significantly diminished CONTRA STMD effect relative to all other groups (Post adults vs. PV+, p < 0.001; Post PV- vs. PV+, p < 0.001; Post juveniles vs. PV+, p < 0.001). **Group definitions:** **Adults:** C57BL/6J adult mice **Juveniles:** C57BL/6J mice at postnatal day 28 **PV-:** adult PV-Cre mice with reduced PV cell activity in V1 **PV+:** adult PV-Cre mice with enhanced PV cell activity in V1 Error bars indicate s.e.m. Statistical significance: *p < 0.05; **p < 0.01; ***p < 0.001.

As in adults, the pronounced reduction in the C/I VEP ratio was driven by a significant decrease in CONTRA VEP amplitudes (Pre = 96.41 ± 12.47 μV vs. Post = 54.56 ± 9.91 μV, Two-Way RM ANOVA, eye × time, DF = 2, F = 75.36; Tukey method, p = 0.001; **Figure 3E,G**) and a concurrent increase in IPSI VEP amplitudes (Pre = 34.83 ± 4.42 μV vs. Post = 70.46 ± 13.70 μV, p < 0.01; **Figure 3F,G**).

The pre-deprivation CONTRA/IPSI asymmetry (Pre Contra = 96.41 ± 12.47 μV vs. Pre Ipsi = 34.83 ± 4.42 μV, p < 0.001) was abolished, and even transiently reversed, after STMD, with IPSI VEP responses exceeding CONTRA responses, although this difference was not statistically significant (Post Contra = 54.56 ± 9.91 μV vs. Post Ipsi = 70.46 ± 13.70 μV, p = 0.270). Normal asymmetry was restored after 2 hours of binocular vision (Contra After 2h = 98.48 ± 12.35 μV; Ipsi After 2h = 36.43 ± 4.66 μV; p < 0.001; **Figure 3D,G**).

#### 2.2 CONTRA eye deprivation in juvenile mice

In a separate group of juvenile mice (n = 8; P28) baseline C/I VEP ratio was 2.83 ± 0.013. CONTRA STMD caused a significant increase in the ratio (Pre = 2.83 ± 0.013 vs. Post = 4.561 ± 0.132, One-Way RM ANOVA, DF = 2, F = 156.1; Tukey method, p < 0.001). This effect was transient, returning to baseline after 2 hours of binocular vision (One-Way RM ANOVA, Pre = 2.83 ± 0.013 vs. After 2h = 2.824 ± 0.013, p = 0.604; **Figure 4C**).

The OD shift after CONTRA STMD in juvenile mice was more pronounced than in adults, as indicated by the higher Post-deprivation C/I VEP ratio (Post adult = 4.080 ± 0.117 vs. Post P28 = 4.561 ± 0.132; t = -2.706, DF = 14.483, two-tailed unpaired t-test, p < 0.05; **Figure 5B**). This change was driven by a significant increase in CONTRA VEP amplitudes (Two-Way RM ANOVA, eye × time, DF = 2, F = 11.08; Tukey method, Pre = 78.98 ± 5.78 μV vs. Post = 99.31 ± 7.68 μV, p < 0.001; **Figure 4E,G**) and a non-significant decrease in IPSI responses (Pre = 28.24 ± 2.13 μV vs. Post = 21.65 ± 1.62 μV, p = 0.689; **Figure 4F,G**). CONTRA dominance over IPSI responses remained significant across all three sessions (Two-Way RM ANOVA, p < 0.001; **Figure 4D,G**).

Together, these results show that STMD induces a rapid and transient OD shift in both adult mice and juvenile animals, indicating that rapid homeostatic plasticity can occur both within and beyond the classical CP. Importantly, the magnitude of the effect is greater in juvenile animals compared to adults.

### 3. Short-term binocular deprivation does not produce any OD shift in adult mice

Binocular deprivation during the CP is known to produce weaker effects than MD (Antonini et al., 1998), likely because the absence of interocular competition reduces the drive for OD shifts (Kasamatsu & Imamura, 2020). Additionally, the OD shift observed after STMD might partly reflect increased retinal sensitivity to light in the previously deprived eye after reopening (Kawasaki et al., 2018). To determine whether this rapid homeostatic plasticity requires interocular competition, and to assess if the observed effects of STMD could be explained solely by enhanced sensitivity of the deprived eye, we applied the same three-session recording protocol (Pre, Post, After 2h; **Figure 6A,B**) to adult mice subjected to binocular deprivation (STBD).

**Figure 6.**
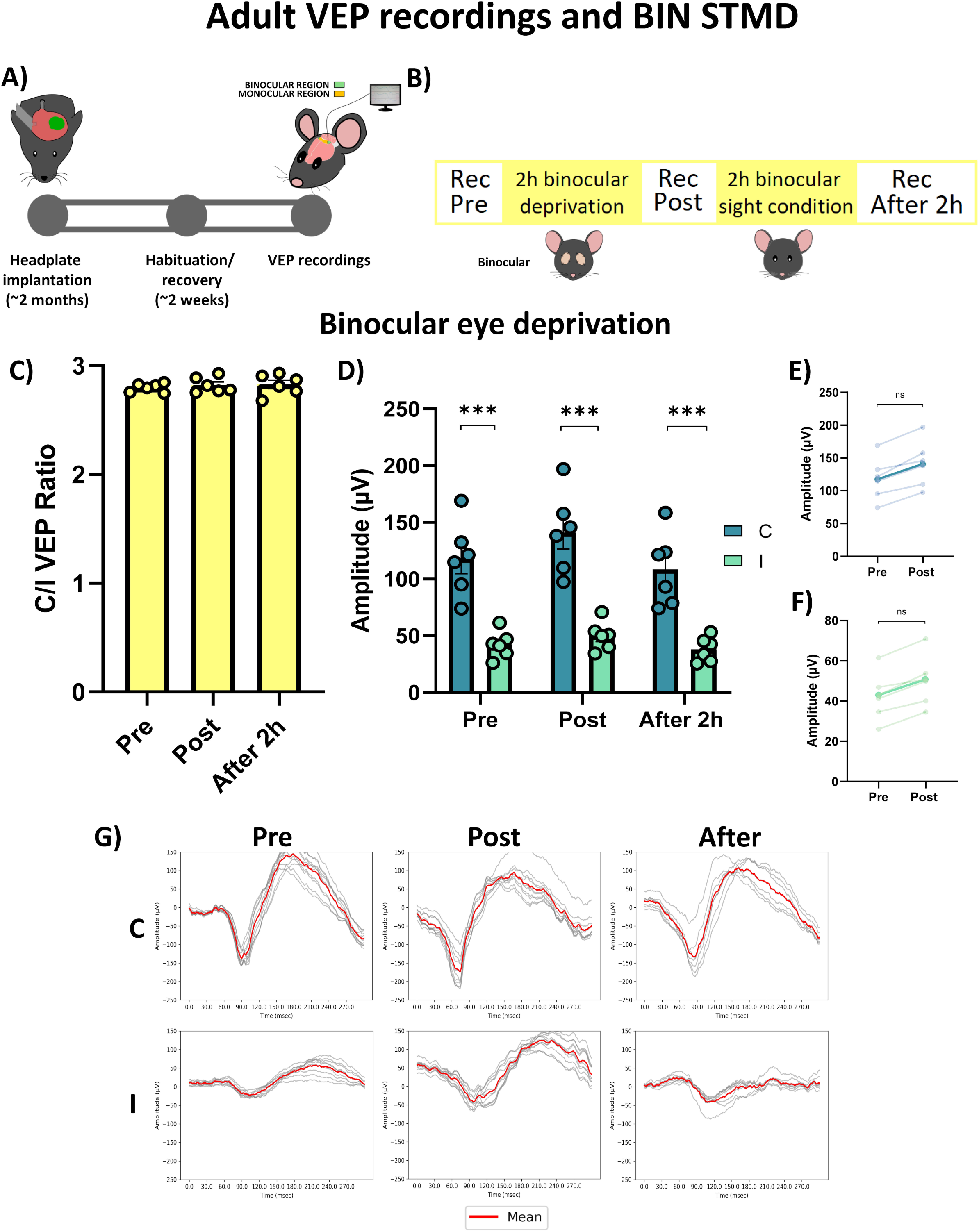
Short-term binocular deprivation (STBD) in adult mice. **a**, Schematic illustration of the experimental protocol for STBD. **b,** Timeline of electrophysiological recordings: baseline recordings before STBD (Pre), recordings immediately after STBD (Post), and recordings following two hours of restored binocular vision (After 2h). **c,** Contralateral/Ipsilateral (C/I) VEP ratio in response to a low spatial frequency grating stimulus (0.06 c/deg). STBD did not produce significant changes in the C/I VEP ratio (One-way ANOVA RM, p=0.503). **d,** Mean VEP amplitudes recorded from contralateral and ipsilateral eyes at all timepoints. **e,** Mean VEP amplitudes from the contralateral eye before and after STBD. Two hours of binocular deprivation results in a non-significant increase in contralateral VEP amplitudes (Two-way ANOVA RM, p=0.072). The darkest line represents the mean. **f,** Mean VEP amplitudes from the ipsilateral eye before and after STBD. Two hours of binocular deprivation induces a non-significant increase in ipsilateral VEP amplitudes (Two-way ANOVA RM, p=0.921). The darkest line represents the mean. **g,** Representative VEP traces from one mouse, demonstrating changes across the different timepoints. C: contralateral eye; I: ipsilateral eye; Pre: before STBD; Post: after STBD; After: two hours after the end of binocular deprivation. Statistical significance: ns: not significant; *p < 0.05; **p < 0.01; ***p < 0.001; error bars indicate S.E.M.

Awake, head-fixed adult mice (n = 6; ∼P74) underwent STBD. Baseline measurements showed a C/I VEP ratio of 2.798 ± 0.016, consistent with normal OD. The C/I VEP ratio remained stable following STBD, both immediately post-deprivation and 2 hours after eye reopening (Pre = 2.798 ± 0.016, Post = 2.822 ± 0.029, After 2h = 2.827 ± 0.038; One-Way ANOVA, DF = 2, F = 0.736; Tukey method, p = 0.503; **Figure 6C**), indicating no shift in OD. STBD did not significantly alter VEP amplitudes for either the CONTRA eye (Pre = 117.866 ± 13.261 μV vs. Post = 141.008 ± 14.469 μV, Two-Way RM ANOVA, eye × time, DF = 2, F = 75.36; Tukey method, p = 0.072; **Figure 6E,G**) or the IPSI eye (Pre = 42.255 ± 4.882 μV vs. Post = 50.02 ± 5.130 μV, p = 0.921; **Figure 6F,G**), confirming the absence of OD changes. Throughout all three recording sessions, the CONTRA eye maintained its dominance over the IPSI eye (Two-Way RM ANOVA, p < 0.001; **Figure 6D,G**), further indicating that STBD did not induce any OD shift. These findings indicate that interocular imbalance is necessary for the rapid homeostatic plasticity observed with STMD, and that the OD shifts cannot be explained solely by enhanced light sensitivity of the previously deprived eye.

### 4. V1 interhemispheric crosstalk modulates OD shifts after short-term monocular deprivation

To assess whether interhemispheric V1 activity contributes to STMD-induced plasticity, we chemogenetically inhibited the contralateral V1 during the 2 h of deprivation. Blocking the contralateral hemisphere partially attenuated the OD shift for both IPSI and CONTRA STMD, but did not abolish it, indicating that interhemispheric interactions modulate, but are not essential for, rapid homeostatic plasticity (see **Supplementary Figs. 1–2** for full VEP amplitudes and statistics).

### 5. Chemogenetic modulation of parvalbumin interneuron activity in V1 during short-term monocular deprivation in adult PV-Cre mice

We used chemogenetics to selectively suppress or enhance PV interneuron activity in V1, aiming to test their causal role in STMD-induced OD plasticity. Adult PV-Cre mice were injected bilaterally in V1 with Cre-dependent AAVs expressing inhibitory (hM4D(Gi)) or excitatory (hM4D(Gq)) DREADDs selectively in PV interneurons. Bilateral injections were performed to mitigate the influence of the non-recorded hemisphere observed in the previous V1 blockade experiments. Immunohistochemistry confirmed PV-specific viral expression restricted to V1 (**Figures 7, 8**).

**Figure 7.**
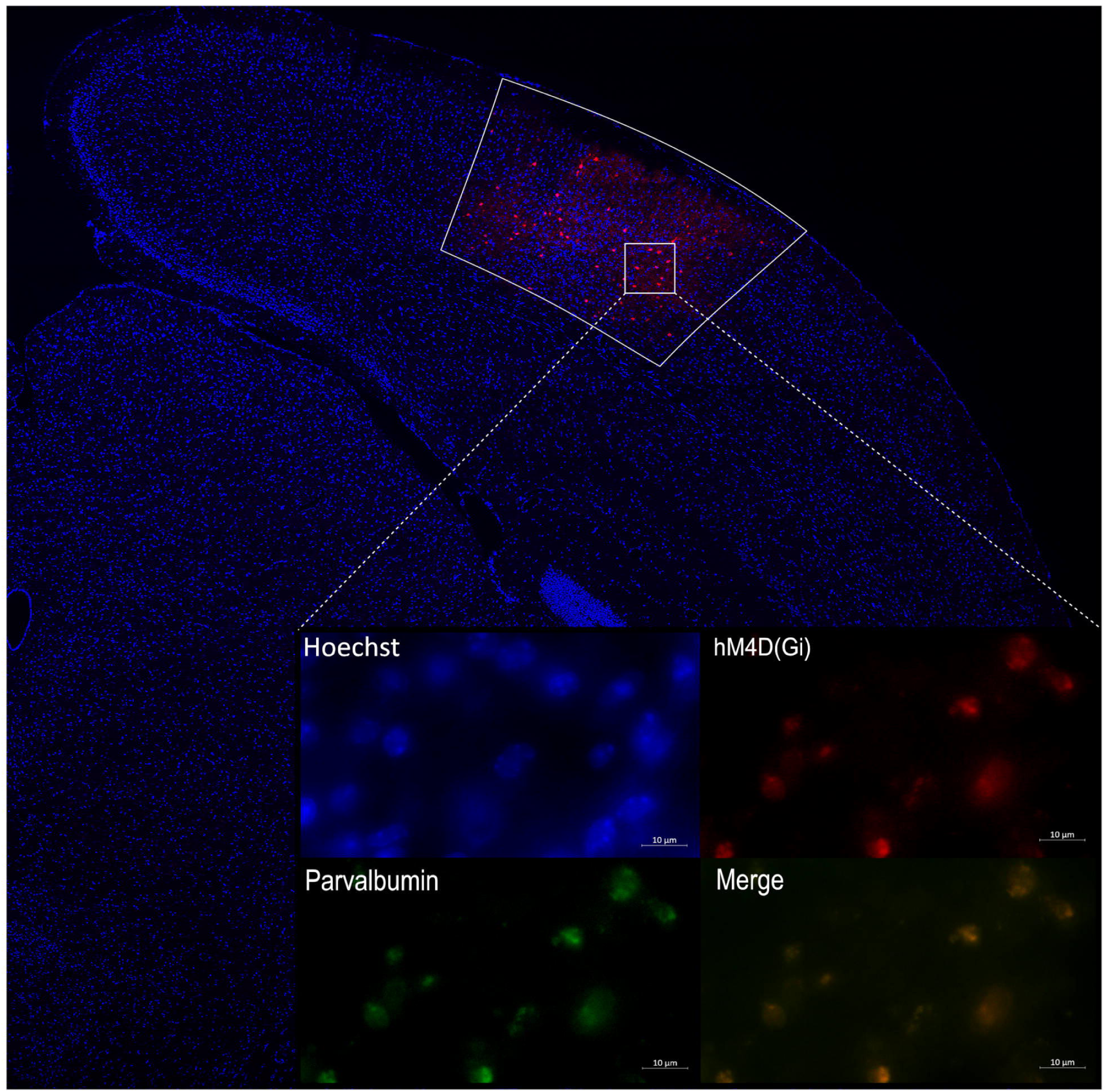
Immunohistochemical analysis of AAV-mediated transgene expression in the primary visual cortex (V1). **a**, Expression of hM4D(Gi) was confined within the white-bordered anatomical boundaries of V1. **b,** Viral expression was specifically localized to parvalbumin-positive interneurons.

**Figure 8.**
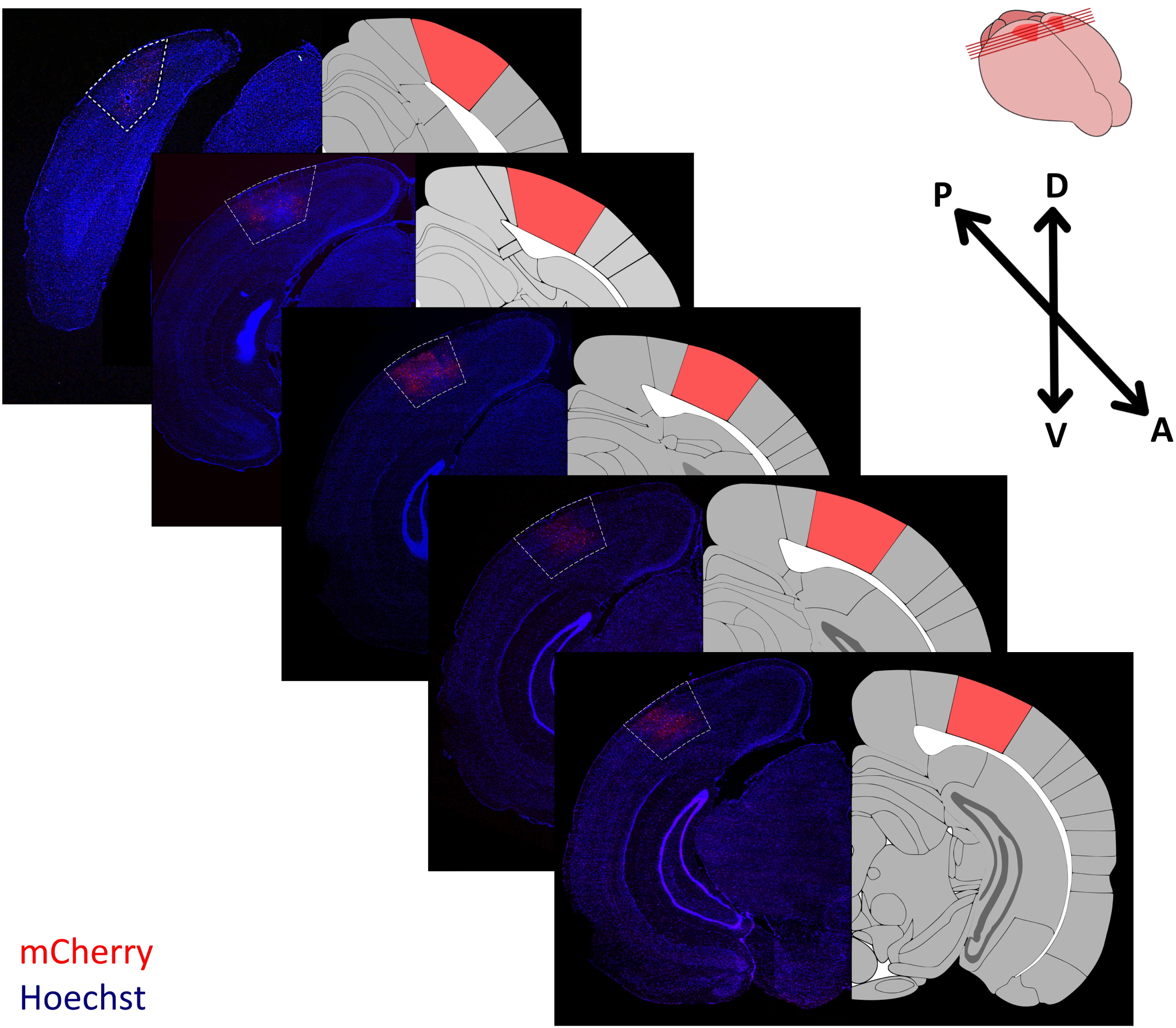
Histological evaluation of mCherry expression. **a**, Histological assessment of mCherry expression was performed to indirectly evaluate hM4D(Gi) expression. **b,** Fluorescent microscopy was used to image 50 μm-thick brain slices along the posterior (P) to anterior (A) axis. **c,** The boundaries of the primary visual cortex (V1) were delineated based on the Allen Brain Atlas. **d,** In the upper right corner, dorsal (D), ventral (V), anterior (A), and posterior (P) coordinates are indicated. **e,** The red lines on the mouse brain reconstruction represent the anatomical positions of the brain slices along the A-P axis.

#### 5.1 Reducing PV interneuron activity in V1 enhances OD plasticity during STMD

To test whether PV inhibition enhances OD plasticity, two groups of awake PV-Cre mice underwent the standard STMD protocol (three VEP recordings: Pre, Post, After 2 h; **Figures 9A,B, 10A,B**), with PV activity suppressed by two hourly intraperitoneal CNO injections during deprivation.

**Figure 9.**
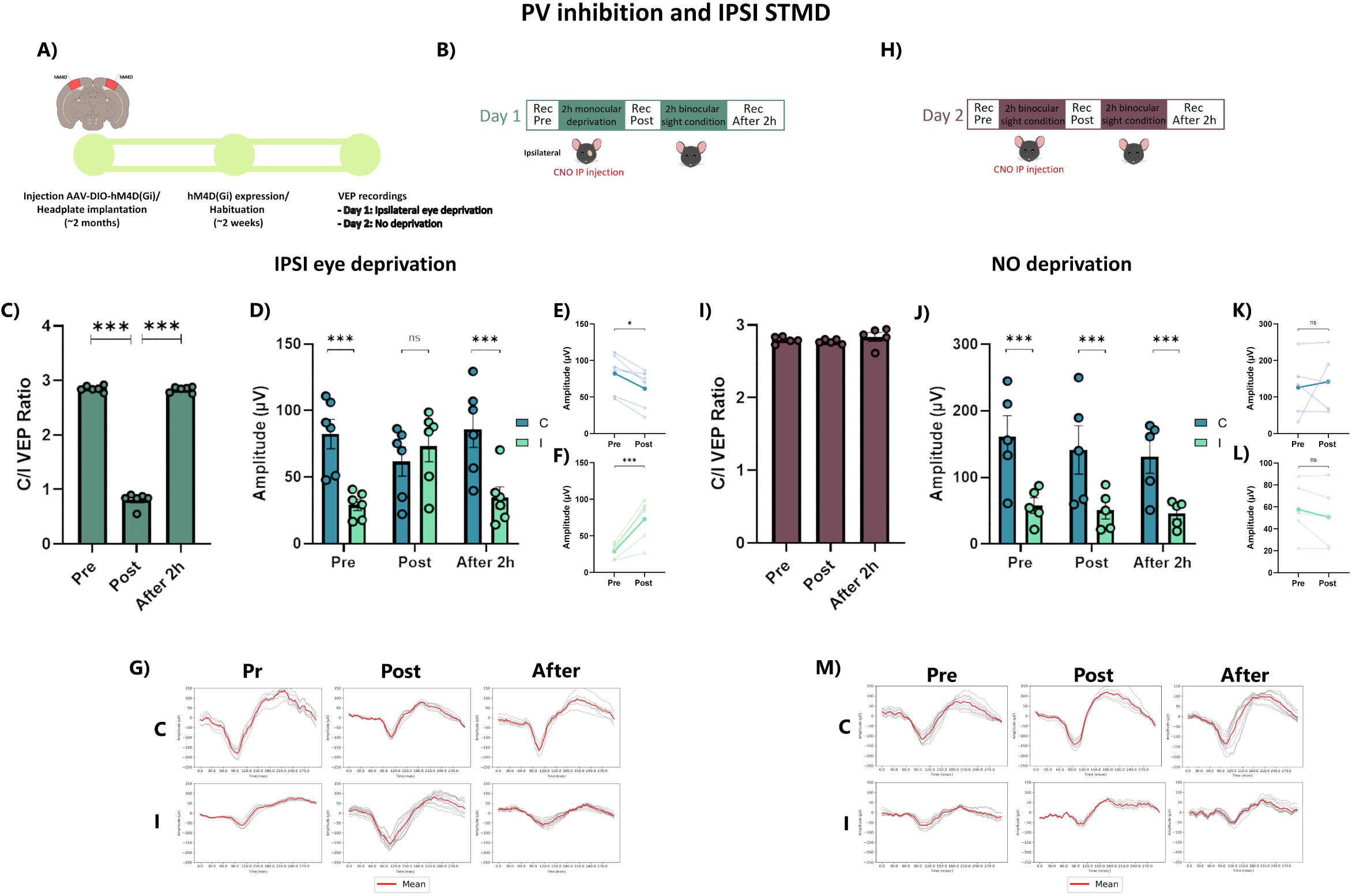
Chemogenetic inhibition of parvalbumin (PV) interneurons activity in V1 during IPSI short-term monocular deprivation (STMD) in adult PV-Cre mice. **a**, Schematic illustration of the experimental protocol. **b,** Timeline of electrophysiological recordings on Day 1: baseline measurements were taken prior to IPSI STMD combined with reduction of PV cells activity in V1 (Pre). Recordings were then conducted immediately after STMD, with PV cells activity restored (Post), followed by recordings performed after two hours of restored binocular vision (After 2h). **c,** Contralateral/Ipsilateral (C/I) VEP ratio in response to a low spatial frequency grating stimulus (0.06 c/deg). The C/I VEP ratio significantly decreased immediately after IPSI STMD compared to baseline (One-way ANOVA RM, p<0.001), and returned to baseline levels following two hours of binocular vision (One-way RM ANOVA, p=0.928). **d,** Mean VEP amplitudes recorded from contralateral and ipsilateral eyes at all timepoints. **e,** Mean VEP amplitudes from the contralateral eye before and after IPSI STMD. Two hours of monocular deprivation resulted in a significant reduction in contralateral VEP amplitudes (Two-way ANOVA RM, p<0.05). The darkest line represents the mean. **f,** Mean VEP amplitudes from the ipsilateral eye before and after IPSI STMD. Two hours of monocular deprivation induced a significant increase in ipsilateral VEP amplitudes (Two-way ANOVA RM, p<0.001). The darkest line represents the mean. **g,** Representative VEP traces from one mouse, demonstrating changes across the different timepoints. **h,** Timeline of electrophysiological recordings on Day 2: baseline measurements (Pre), after two hours of reduced PV interneuron activity in V1 without monocular deprivation (Post), and an additional recording obtained after two hours of restored PV activity in V1 (After 2h). **i,** Three consecutive electrophysiological recordings with two hours of reduction of PV cells activity in V1 without monocular deprivation did not result in significant changes in the C/I VEP ratio across timepoints (One-way ANOVA RM, p=0.451). **j,** Mean VEP amplitudes recorded from contralateral and ipsilateral eyes at all timepoints. **k,** Mean contralateral eye VEP amplitudes at baseline (Pre) and after two hours of binocular vision combined with PV cells activity reduction in V1 (Post). No significant differences were observed between Pre- and Post-recordings (Two-way ANOVA RM, p=0.893). The darkest line represents the mean. **l,** Mean ipsilateral eye VEP amplitudes at baseline (Pre) and after two hours of binocular vision combined with PV cells activity reduction in V1 (Post). No significant differences were observed between Pre- and Post-recordings (Two-way ANOVA RM, p=0.998). The darkest line represents the mean. **m,** Representative VEP traces from one mouse across the different timepoints. C: contralateral eye; I: ipsilateral eye; Pre: before STMD; Post: after STMD; After: two hours after the end of monocular deprivation. Statistical significance: ns: not significant; *p < 0.05; **p < 0.01; **p < 0.001; error bars indicate S.E.M.

##### 5.1.1 PV interneuron inhibition during IPSI eye short-term deprivation

In PV-Cre mice (n=6), baseline C/I VEP ratio was 2.855 ± 0.022. IPSI STMD with PV inhibition caused a marked decrease (Post=0.810 ± 0.052, One-way ANOVA RM; DF=2, F=1022; Tukey method, p<0.001), which returned to baseline after 2 h of binocular vision (2.836 ± 0.021; One-way ANOVA RM, p=0.928; **Figure 9C**). This shift reflected both a decrease in CONTRA VEP amplitude (Pre=82.40±11.01 μV; Post=61.78±10.83 μV; Two-way ANOVA RM; eye x time, DF=2, F=41.26, Tukey method, p<0.05; **Figure 9E,G**) and an increase in IPSI responses (Pre=29.17±4.09 μV, Post=73.20±11.54 μV; Two-way ANOVA RM, p<0.001; **Figure 9F,G**). Pre-deprivation contralateral dominance (Two-way ANOVA RM, p<0.001) was reversed post-STMD and restored after 2 h (Two-way ANOVA RM, p<0.001; **Figure 9D,G**). The OD shift in PV-Cre mice was larger than in adult wild-types (two-tailed unpaired t-test, Post WT=1.101±0.093 vs. Post PV^-^=0.810±0.052; t=2.725, DF=9.2395; p<0.05) and comparable to P28 juveniles (two-tailed unpaired t-test, Post PV^-^=0.810±0.052 vs. Post P28=0.819±0.033; t=-0.144, DF=8.9789; p=0.888; **Figure 5A**). CNO alone (no MD) had no effect on C/I ratio or VEP amplitudes (**Figure 9H–M**).

##### 5.1.2 PV interneuron inhibition during CONTRA eye short-term deprivation

In another group (n=6), CONTRA STMD with PV inhibition significantly increased the C/I ratio (Pre=2.891±0.024 vs. Post=4.613±0.02; One-way ANOVA RM; DF=2, F=1071; Tukey method, p<0.001), which returned to baseline after 2 h of binocular vision (After 2h=2.865±0.03, p=0.821; One-way ANOVA RM, p=0.821; **Figure 10C**). The increase was driven by enhanced CONTRA responses (Pre=115.586±5.697 LμV; Post=156.383±15.052 μV; Two-way ANOVA RM; eye x time, DF=2, F=382.97, Tukey method, p<0.001; **Figure 10E–G**), while IPSI responses showed a small, non-significant decrease (Pre=40.121±2.197 μV, Post=34.043±3.242 μV; Two-way ANOVA RM, p=0.963; **Figure 10F–G**). This OD shift exceeded that of wild-type adults (two-tailed unpaired t-test, Post WT=4.080±0.117 vs. Post PV^-^=4.613±0.02; t=-4.462, DF=8.491; p<0.01) and matched P28 juveniles (two-tailed unpaired t-test, Post PV^-^=4.613±0.02 vs. Post P28=4.561±0.132; t=0.390, DF=7.338; p=0.707; **Figure 5B**). CNO without MD produced no changes in C/I ratio or VEP amplitudes (**Figure 10 H–M**).

**Figure 10.**
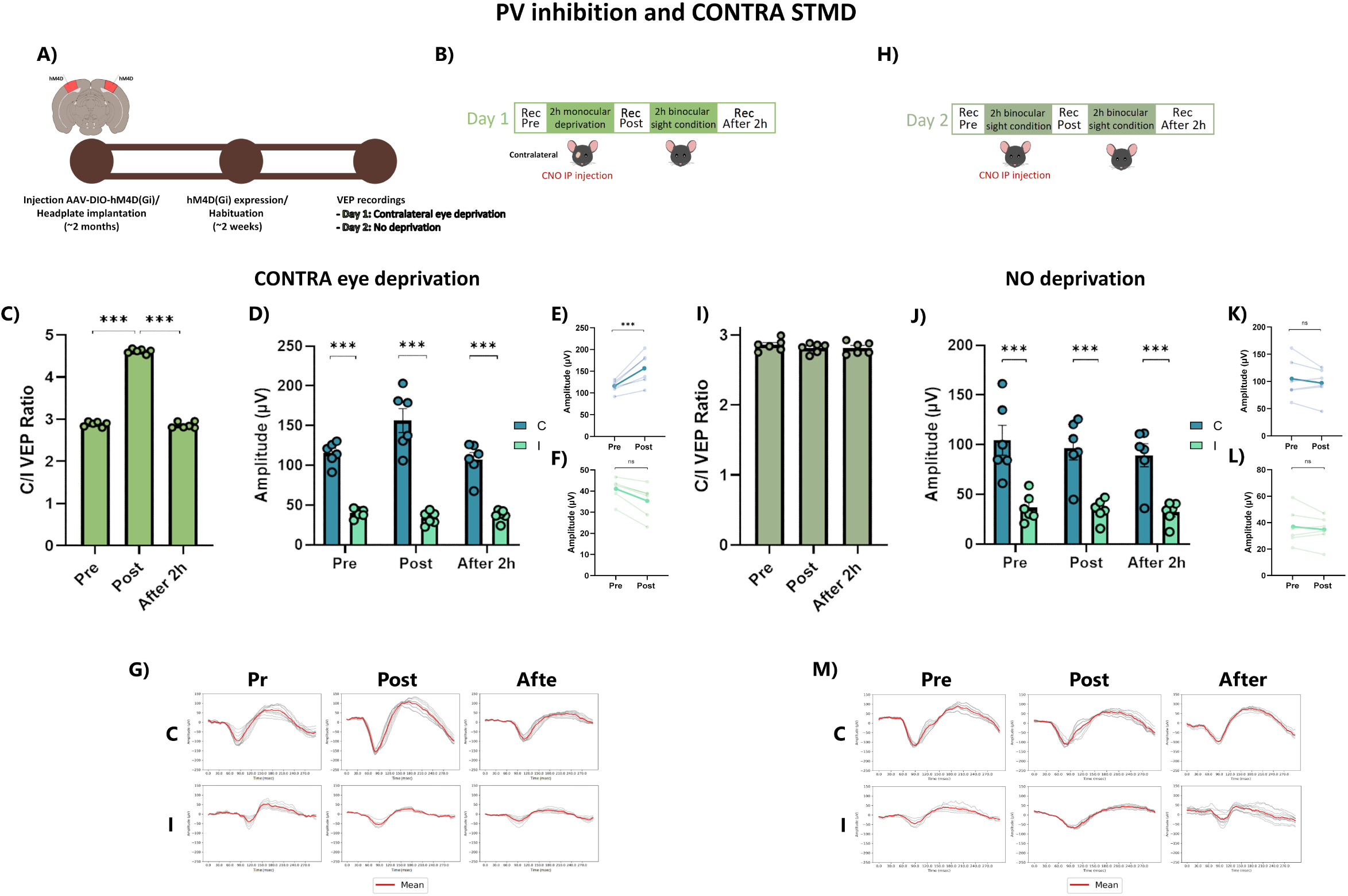
Chemogenetic inhibition of parvalbumin (PV) interneurons activity in V1 during CONTRA short-term monocular deprivation (STMD) in adult PV-Cre mice. **a**, Schematic illustration of the experimental protocol. **b,** Timeline of electrophysiological recordings on Day 1: baseline measurements were taken prior to CONTRA STMD combined with reduction of PV cells activity in V1 (Pre). Recordings were then conducted immediately after STMD, with PV cells activity restored (Post), followed by recordings performed after two hours of restored binocular vision (After 2h). **c,** Contralateral/Ipsilateral (C/I) VEP ratio in response to a low spatial frequency grating stimulus (0.06 c/deg). The C/I VEP ratio significantly increased immediately after CONTRA STMD compared to baseline (One-way ANOVA RM, p<0.001), and returned to baseline levels following two hours of binocular vision (One-way RM ANOVA, p=0.821). **d,** Mean VEP amplitudes recorded from contralateral and ipsilateral eyes at all timepoints. **e,** Mean VEP amplitudes from the contralateral eye before and after CONTRA STMD. Two hours of monocular deprivation resulted in a significant increase in contralateral VEP amplitudes (Two-way ANOVA RM, p<0.001). The darkest line represents the mean. **f,** Mean VEP amplitudes from the ipsilateral eye before and after CONTRA STMD. Two hours of monocular deprivation induced a non-significant reduction in ipsilateral VEP amplitudes (Two-way ANOVA RM, p=0.963). The darkest line represents the mean. **g,** Representative VEP traces from one mouse, demonstrating changes across the different timepoints. **h,** Timeline of electrophysiological recordings on Day 2: baseline measurements (Pre), after two hours of reduced PV interneuron activity in V1 without monocular deprivation (Post), and an additional recording obtained after two hours of restored PV activity in V1 (After 2h). **i,** Three consecutive electrophysiological recordings with two hours of reduction of PV cells activity in V1 without monocular deprivation did not result in significant changes in the C/I VEP ratio across timepoints (One-way ANOVA RM, p=0.149). **j,** Mean VEP amplitudes recorded from contralateral and ipsilateral eyes at all timepoints. **k,** Mean contralateral eye VEP amplitudes at baseline (Pre) and after two hours of binocular vision combined with PV cells activity reduction in V1 (Post). No significant differences were observed between Pre- and Post-recordings (Two-way ANOVA RM, p=0.937). The darkest line represents the mean. **l,** Mean ipsilateral eye VEP amplitudes at baseline (Pre) and after two hours of binocular vision combined with PV cells activity reduction in V1 (Post). No significant differences were observed between Pre- and Post-recordings (Two-way ANOVA RM, p=0.999). The darkest line represents the mean. **m,** Representative VEP traces from one mouse across the different timepoints. C: contralateral eye; I: ipsilateral eye; Pre: before STMD; Post: after STMD; After: two hours after the end of monocular deprivation. Statistical significance: ns: not significant; *p < 0.05; **p < 0.01; **p < 0.001; error bars indicate S.E.M.

#### 5.2 Enhancing PV interneuron activity blocks OD plasticity during short-term monocular deprivation

To determine whether increasing PV interneuron activity limits STMD-induced plasticity, we repeated the protocol with excitatory DREADDs (hM4D(Gq)) in V1 (**Figure 11A,B**; **Figure 12A,B**), administering CNO as in previous experiments.

**Figure 11.**
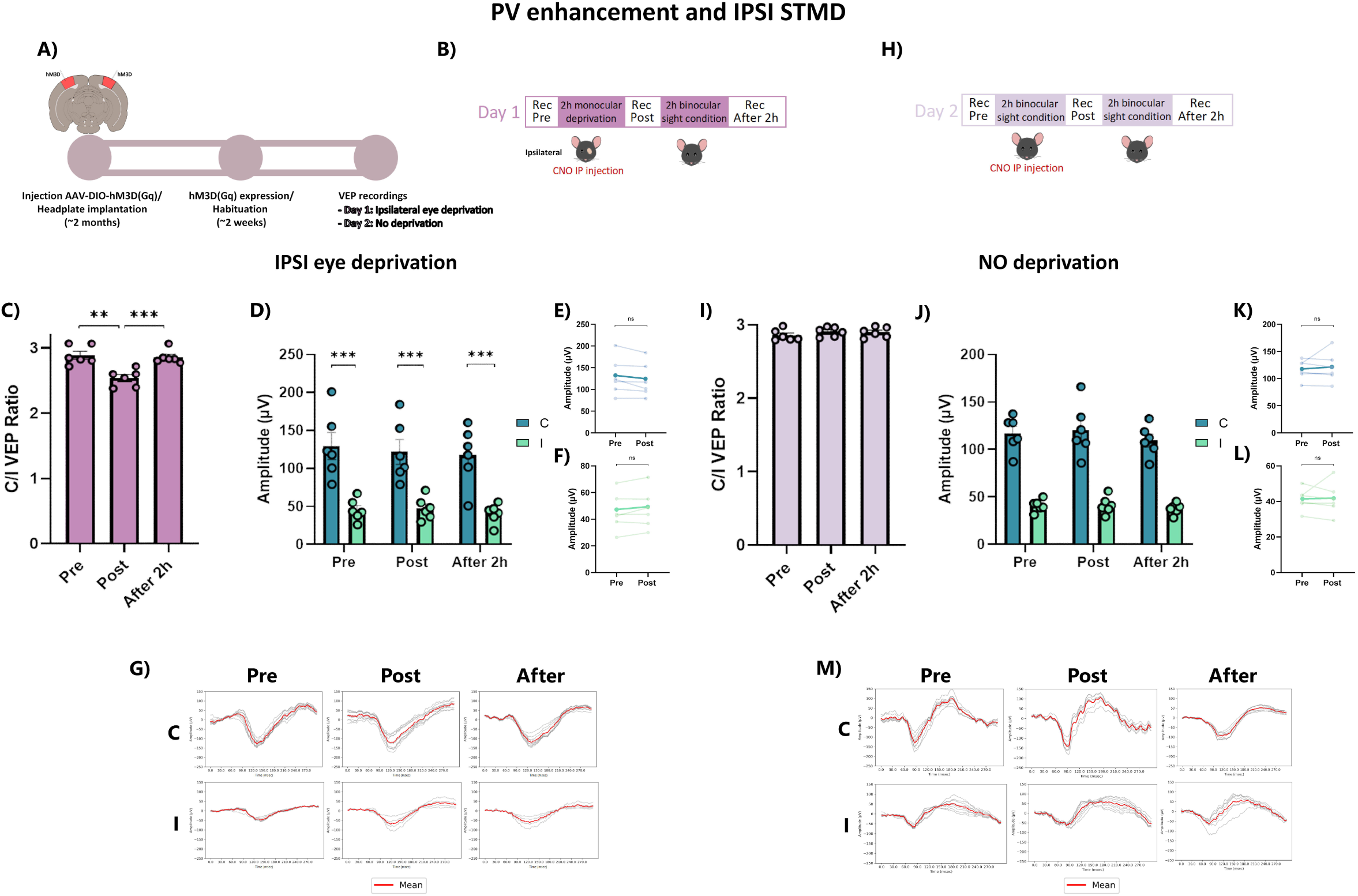
Chemogenetic enhancement of parvalbumin (PV) interneurons activity in V1 during IPSI short-term monocular deprivation (STMD) in adult PV-Cre mice. **a**, Schematic illustration of the experimental protocol. **b,** Timeline of electrophysiological recordings on Day 1: baseline measurements were taken prior to IPSI STMD combined with enhancement of PV cells activity in V1 (Pre). Recordings were then conducted immediately after STMD, with PV cells activity restored (Post), followed by recordings performed after two hours of restored binocular vision (After 2h). **c,** Contralateral/Ipsilateral (C/I) VEP ratio in response to a low spatial frequency grating stimulus (0.06 c/deg). The C/I VEP ratio significantly decreased immediately after IPSI STMD compared to baseline (One-way ANOVA RM, p<0.01), and returned to baseline levels following two hours of binocular vision (One-way RM ANOVA, p=0.902). **d,** Mean VEP amplitudes recorded from contralateral and ipsilateral eyes at all timepoints. **e,** Mean VEP amplitudes from the contralateral eye before and after IPSI STMD. Two hours of monocular deprivation resulted in a non-significant reduction in contralateral VEP amplitudes (Two-way ANOVA RM, p=0.962). The darkest line represents the mean. **f,** Mean VEP amplitudes from the ipsilateral eye before and after IPSI STMD. Two hours of monocular deprivation induced a non-significant decrease in ipsilateral VEP amplitudes (Two-way ANOVA RM, p=0.999). The darkest line represents the mean. **g,** Representative VEP traces from one mouse, demonstrating changes across the different timepoints. **h,** Timeline of electrophysiological recordings on Day 2: baseline measurements (Pre), after two hours of enhanced PV interneuron activity in V1 without monocular deprivation (Post), and an additional recording obtained after two hours of restored PV activity in V1 (After 2h). **i,** Three consecutive electrophysiological recordings with two hours of enhancement of PV cells activity in V1 without monocular deprivation did not result in significant changes in the C/I VEP ratio across timepoints (One-way ANOVA RM, p=0.284). **j,** Mean VEP amplitudes recorded from contralateral and ipsilateral eyes at all timepoints. **k,** Mean contralateral eye VEP amplitudes at baseline (Pre) and after two hours of binocular vision combined with PV cells activity enhancement in V1 (Post). No significant differences were observed between Pre- and Post-recordings (Two-way ANOVA RM, p=0.995). The darkest line represents the mean. **l,** Mean ipsilateral eye VEP amplitudes at baseline (Pre) and after two hours of binocular vision combined with PV cells activity enhancement in V1 (Post). No significant differences were observed between Pre- and Post-recordings (Two-way ANOVA RM, p=1.00). The darkest line represents the mean. **m,** Representative VEP traces from one mouse across the different timepoints. C: contralateral eye; I: ipsilateral eye; Pre: before STMD; Post: after STMD; After: two hours after the end of monocular deprivation. Statistical significance: ns: not significant; *p < 0.05; **p < 0.01; **p < 0.001; error bars indicate S.E.M.

**Figure 12.**
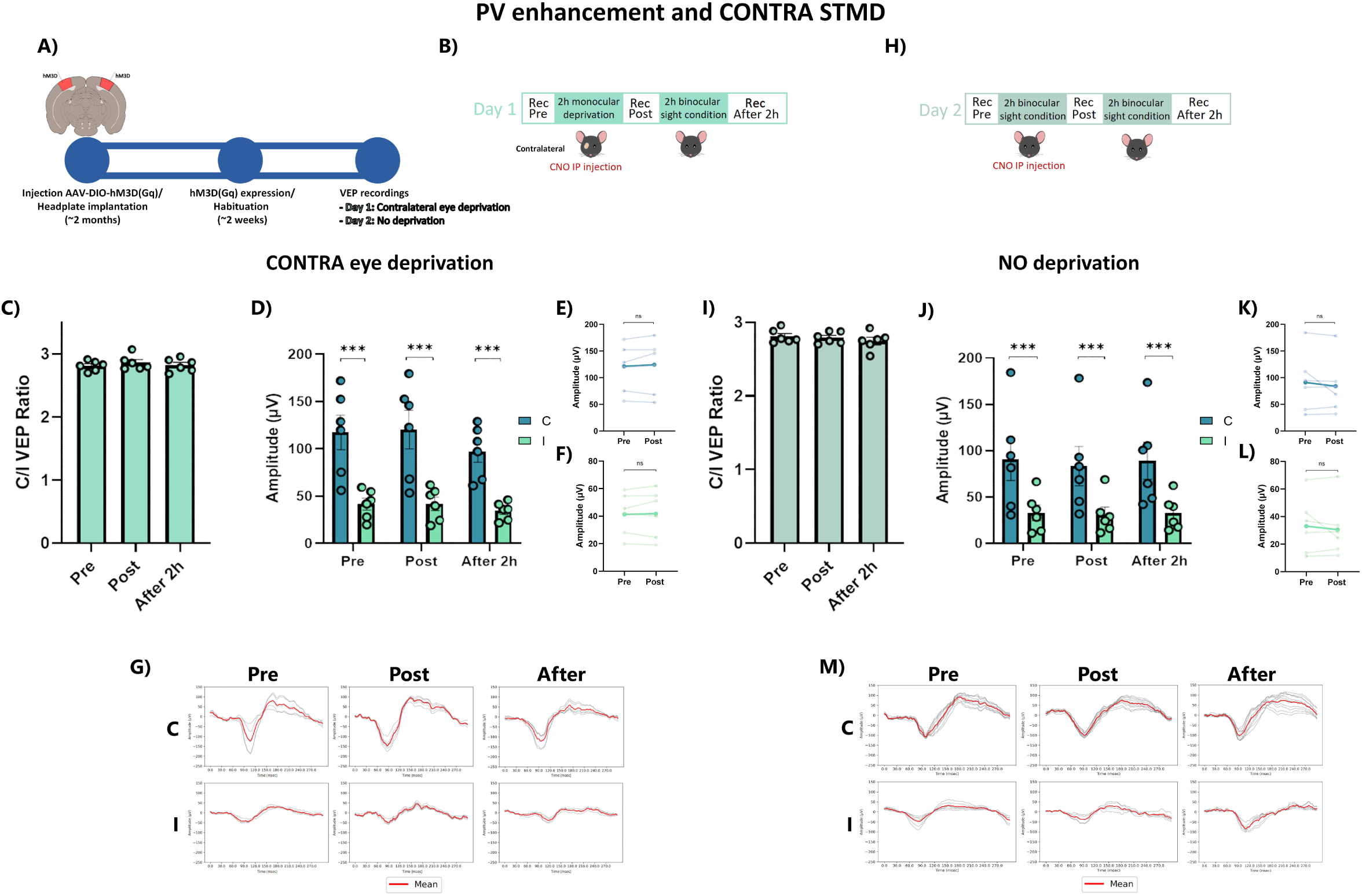
Chemogenetic enhancement of parvalbumin (PV) interneurons activity in V1 during CONTRA short-term monocular deprivation (STMD) in adult PV-Cre mice. **a**, Schematic illustration of the experimental protocol. **b,** Timeline of electrophysiological recordings on Day 1: baseline measurements were taken prior to CONTRA STMD combined with enhancement of PV cells activity in V1 (Pre). Recordings were then conducted immediately after STMD, with PV cells activity restored (Post), followed by recordings performed after two hours of restored binocular vision (After 2h). **c,** The increase in PV cell activity in V1 during CONTRA STMD completely abolished the OD shift, as evidenced by no significant changes in the C/I VEP ratio across timepoints (One-way RM ANOVA, p=0.280). **d,** Mean VEP amplitudes recorded from contralateral and ipsilateral eyes at all timepoints. **e,** Mean VEP amplitudes from the contralateral eye before and after CONTRA STMD. When PV cell activity is increased, two hours of monocular deprivation do not induce changes in contralateral VEP amplitudes (Two-way ANOVA RM, p=0.999). The darkest line represents the mean. **f,** Mean VEP amplitudes from the ipsilateral eye before and after CONTRA STMD. Consistent with observations in contralateral VEP amplitudes, increasing PV cells activity prevents changes in ipsilateral VEP amplitudes following two hours of monocular deprivation (Two-way ANOVA RM, p=1.00). The darkest line represents the mean. **g,** Representative VEP traces from one mouse, demonstrating changes across the different timepoints. **h,** Timeline of electrophysiological recordings on Day 2: baseline measurements (Pre), after two hours of enhanced PV interneuron activity in V1 without monocular deprivation (Post), and an additional recording obtained after two hours of restored PV activity in V1 (After 2h). **i,** Three consecutive electrophysiological recordings with two hours of enhancement of PV cells activity in V1 without monocular deprivation did not result in significant changes in the C/I VEP ratio across timepoints (One-way ANOVA RM, p=0.250). **j,** Mean VEP amplitudes recorded from contralateral and ipsilateral eyes at all timepoints. **k,** Mean contralateral eye VEP amplitudes at baseline (Pre) and after two hours of binocular vision combined with PV cells activity enhancement in V1 (Post). No significant differences were observed between Pre- and Post-recordings (Two-way ANOVA RM, p=0.983). The darkest line represents the mean. **l,** Mean ipsilateral eye VEP amplitudes at baseline (Pre) and after two hours of binocular vision combined with PV cells activity enhancement in V1 (Post). No significant differences were observed between Pre- and Post-recordings (Two-way ANOVA RM, p=0.999). The darkest line represents the mean. **m,** Representative VEP traces from one mouse across the different timepoints. C: contralateral eye; I: ipsilateral eye; Pre: before STMD; Post: after STMD; After: two hours after the end of monocular deprivation. Statistical significance: ns: not significant; *p < 0.05; **p < 0.01; **p < 0.001; error bars indicate S.E.M.

##### 5.2.1 PV enhancement combined with IPSI short-term deprivation

In PV-Cre mice (n=6), baseline C/I ratio was 2.891 ± 0.059. IPSI STMD combined with PV activation produced a modest reduction in the C/I ratio (Pre=2.891±0.059 vs. Post=2.542±0.053; One-way RM ANOVA; DF=2, F=17.86; Tukey method, p<0.01), which returned to baseline after 2 hours of binocular vision (Pre=2.891±0.059 vs. After 2h=2.863±0.042; One-way RM ANOVA, p=0.902; **Figure 11C**).

CONTRA and IPSI VEP amplitudes remained unchanged (Two-way RM ANOVA; eye × time, DF=2, F=0.575; Tukey method: CONTRA p=0.962, IPSI p=0.999; **Figure 11E-G**), and contralateral dominance persisted (Two-way RM ANOVA; p<0.001; **Figure 11D,G**).

The resulting OD shift was significantly smaller than in wild-type adults (two-tailed unpaired t-test, Post WT=4.080±0.117 vs. Post PV+=2.862±0.047; t=9.601, DF=10.345; p<0.001), P28 juveniles (Post PV+=2.862±0.047 vs. Post P28=4.561±0.132; t=12.039, DF=8.675; p<0.001), and PV-inhibited mice (Post PV-=4.613±0.02 vs. Post PV+=2.862±0.047; t=34.025, DF=6.865; p<0.001; **Figure 5A**). CNO-only controls produced no changes (**Figure 11H-M**).

##### 5.2.2 PV enhancement combined with CONTRA short-term deprivation

In a separate group of PV-Cre mice (n=6), baseline C/I ratio was 2.813 ± 0.032. CONTRA STMD with PV activation did not produce a significant change in the C/I ratio (Pre=2.813±0.032 vs. Post=2.862±0.047; One-way RM ANOVA; DF=2, F=1.448; Tukey, p=0.28), and values remained at baseline after 2 hours of binocular vision (Pre=2.813±0.032 vs. After 2h=2.824±0.042; One-way ANOVA RM, p=0.925; **Figure 12C**).

CONTRA and IPSI VEP amplitudes were unchanged (Two-way RM ANOVA; eye × time, DF=2, F=0.575; CONTRA Pre=117.383±18.11 μV, Post=120.436±20.303 μV, p=0.999; IPSI Pre=41.55±6.194 μV, Post=41.94±6.194 μV, p=1.000 in all comparison; **Figure 12E-G**), and contralateral dominance persisted (Two-way RM ANOVA; p<0.001; **Figure 12D,G**).

Overall, PV activation abolished OD plasticity, yielding significantly smaller shifts than in wild-type adults (two-tailed unpaired t-test, Post WT=4.080±0.117 vs. Post PV+=2.862±0.047; t=9.601, DF=10.345; p<0.001), P28 juveniles (Post PV+=2.862±0.047 vs. Post P28=4.561±0.132; t=12.039, DF=8.675; p<0.001), or PV-inhibited mice (Post PV-=4.613±0.02 vs. Post PV+=2.862±0.047; t=34.025, DF=6.865; p<0.001; **Figure 5B**). CNO-only controls again showed no effect (**Figure 12H-M**). These results show that PV interneurons bidirectionally gate rapid OD plasticity in adult V1, with inhibition suppressing and disinhibition enhancing experience-dependent shifts. Together, these results demonstrate that PV-mediated inhibition acts as a fast, bidirectional gate for short-term, experience-dependent ocular dominance plasticity in adult V1. Transient disinhibition is sufficient to reinstate juvenile-like levels of plasticity, whereas increased PV activity constrains or abolishes these rapid OD shifts, identifying inhibitory control as a key circuit mechanism enabling homeostatic plasticity beyond the critical period.

## Discussion

In humans, STMD transiently enhances the perceptual dominance of the deprived eye, as demonstrated using binocular rivalry tasks (Lunghi et al., 2011). When distinct images are presented simultaneously to the two eyes, they compete for perceptual dominance, resulting in spontaneous alternations between the two images (Tong et al., 2006). The relative dominance duration reflects the functional weight of each eye: in typical individuals, it is approximately balanced, whereas in amblyopes the non-amblyopic eye dominates. Following brief deprivation, however, the previously deprived eye transiently dominates perception for at least two hours (Lunghi et al., 2011, 2013), a phenomenon interpreted as a form of homeostatic plasticity in the adult visual cortex, in which reduced activity in the deprived pathway leads to a compensatory increase in sensitivity or gain (Lunghi, Berchicci, et al., 2015).

Importantly, this plasticity is further enhanced when deprivation is combined with moderate physical activity (Lunghi & Sale, 2015), and is modulated by alignment between visual input and motor behavior, highlighting a role for sensorimotor integration in adult cortical plasticity (Steinwurzel et al., 2023).

Despite increasing evidence in humans, the cellular and circuit-level mechanisms of this plasticity remain poorly understood due to the scarcity of animal studies. To address this, we adapted the human STMD paradigm to mice. In primates, including humans, roughly 50% of retinal projections from each eye reach both hemispheres (Petros et al., 2008), making deprivation of either eye functionally equivalent; hence most human studies select the perceptually dominant eye for deprivation (Binda & Lunghi, 2017; Lunghi, Berchicci, et al., 2015; Lunghi et al., 2011, 2013; Prosper et al., 2023). In contrast, rodents display pronounced anatomical asymmetry: over 90% of retinal ganglion cells project contralaterally, with only ∼3–5% projecting ipsilaterally (Petros et al., 2008). Accordingly, the normal C/I VEP ratio is ∼2.7–3 (Kaplan et al., 2016; Mazziotti et al., 2017; Porciatti et al., 1999). We thus tested whether this baseline asymmetry influences the effects of MD.

Here, we show that deprivation of the IPSI eye produced a robust OD shift favoring the deprived eye, reflected by a significant reduction in the C/I VEP ratio. This effect resulted from both decreased contralateral and increased ipsilateral VEP amplitudes and reversed after two hours of binocular vision. Similar VEP changes are observed in humans, where brief deprivation alters the early C1 component: amplitude increases for the deprived eye and decreases for the non-deprived eye (Lunghi, Berchicci, et al., 2015).

CONTRA STMD also induced significant changes: VEP amplitudes increased in the deprived eye and decreased in the non-deprived eye. Unlike IPSI deprivation, the OD shift here primarily reflected strengthening of the deprived contralateral eye rather than weakening of the ipsilateral non-deprived eye, likely due to the baseline asymmetry: ipsilateral VEPs are approximately threefold lower than contralateral responses, limiting the magnitude of detectable reductions.

Importantly, both paradigms produced OD shifts favoring the deprived eye, while control conditions with repeated recordings over two-hour intervals yielded no significant changes.

We next examined STMD-induced plasticity during the critical period (P28). Using identical IPSI and CONTRA deprivation protocols, juvenile mice displayed more pronounced OD shifts than adults. Notably, during IPSI STMD, ipsilateral VEP amplitudes surpassed contralateral responses, a phenomenon absent in adults, where contralateral VEPs remained dominant despite increases in ipsilateral responses. Similarly, CONTRA STMD produced larger OD shifts in juveniles. These results indicate that while STMD triggers homeostatic plasticity across ages, its magnitude is greater during the CP. Importantly, the qualitative nature of the OD shift was consistent across age groups, suggesting conserved underlying mechanisms. This finding contrasts sharply with classical long-term MD, where age strongly constrains plasticity. For instance, 4 days of MD leads to OD shift in juveniles but not in adults (Gordon et al., 1996). Even longer deprivations in young adults induce only smaller OD shifts (Hofer et al., 2006; Sawtell et al., 2003), reflecting the well-documented age-related decline in OD plasticity (Lehmann & Löwel, 2008). Thus, the novel observation that STMD induces OD plasticity in adults—with a similar time course but reduced magnitude compared to juveniles—is particularly intriguing. These observations raise the question of how STMD-induced plasticity relates to the classical homeostatic mechanisms engaged by prolonged MD during the CP. Extensive prior work has shown that long-term deprivation recruits a combination of Hebbian competition and homeostatic processes that stabilize cortical activity, including synaptic scaling, changes in intrinsic excitability, and adjustments in inhibitory tone (Kaneko et al., 2008; Mrsic-Flogel et al., 2007; Ranson et al., 2012; G. Turrigiano et al., 1998; G. G. Turrigiano & Nelson, 2004). In the context of ocular dominance plasticity, these mechanisms are most evident during the later phases of deprivation, when responses to the open or previously deprived eye recover or increase despite continued sensory imbalance(Barnes et al., 2015; Sato & Stryker, 2008; G. G. Turrigiano, 2017).

A key difference between these classical forms of homeostatic plasticity and the effects observed here lies in their temporal dynamics. Whereas homeostatic adaptations during the critical period typically emerge over days of deprivation and depend on molecular pathways such as TNFα signaling and synaptic scaling of excitatory synapses (Kaneko et al., 2008; G. G. Turrigiano, 2017), STMD-induced plasticity develops within hours, is rapidly reversible, and is robustly expressed in adult animals.

Importantly, however, these differences do not necessarily imply that STMD engages an entirely distinct homeostatic mechanism. Instead, STMD may reflect an early, fast-acting phase of homeostatic regulation that precedes—and potentially triggers—the slower synaptic and molecular adaptations observed after prolonged deprivation. In this view, rapid modulation of inhibitory circuits, particularly via parvalbumin-positive interneurons, could provide a flexible and reversible means of adjusting cortical gain on short timescales, while longer deprivation durations recruit additional, more stable forms of synaptic homeostasis. Consistent with this interpretation, we find that transient manipulation of PV interneuron activity bidirectionally gates STMD-induced ocular dominance shifts, supporting a role for inhibition as a rapid control point for experience-dependent gain regulation. Determining whether and how this fast inhibitory mechanism transitions into classical homeostatic processes will require systematic manipulation of deprivation duration, spanning short to prolonged timescales. Such experiments will be necessary to establish whether STMD-induced plasticity represents a distinct form of homeostasis or the initial phase of a continuum of homeostatic adaptations.

Notably, although STMD-induced plasticity is present both during and after the CP, its magnitude is enhanced in juvenile mice, indicating that developmental state modulates the expression of rapid homeostatic responses. Together, these findings position STMD-induced ocular dominance plasticity as a rapid, reversible component of cortical homeostasis that complements the slower, well-characterized mechanisms engaged by prolonged deprivation during development.

To assess whether OD changes require interocular competition, we performed two-hour binocular deprivation (STBD). While OD ratios remained unchanged, VEP amplitudes increased non-significantly for both eyes, suggesting a rapid homeostatic effect of smaller magnitude. This may partially reflect heightened light sensitivity from prior binocular deprivation, though an input imbalance between the eyes, instantiated by MD, appears necessary to produce OD shifts.

To evaluate V1 interhemispheric crosstalk influence, we blocked contralateral V1 (with respect to the recorded hemisphere) during STMD. IPSI and CONTRA STMD still produced OD shifts comparable to naïve animals, whereas V1 blockade without deprivation caused no changes, indicating that STMD plasticity is largely intrahemispheric. Nonetheless, OD shifts were slightly smaller, consistent with some contribution from the contralateral hemisphere in awake animals. Interestingly, interhemispheric crosstalk exerts opposite effects depending on deprivation duration. In adults, blocking the influence of the opposite hemisphere during STMD reduces the magnitude of the OD shift toward the deprived eye. In contrast, during classic long-term MD in juveniles, it inhibits deprived-eye inputs (Pietrasanta et al., 2012; Restani et al., 2009), contributing to the classic effect of strengthening the non-deprived eye and weakening the deprived eye. Indeed, continuously blocking callosal input throughout long-term MD prevents the loss of deprived-eye responsiveness(Restani et al., 2009).Further studies are needed to determine whether interhemispheric crosstalk during STMD increases inhibition of non-deprived eye inputs or, conversely, enhances deprived eye inputs.

Excitatory/inhibitory (E/I) balance is a key regulator of OD plasticity (Chen et al., 2022; Fagiolini et al., 2004; Fagiolini & Hensch, 2000; Hensch, 2003, 2005, 2018; Hensch et al., 1998; Maya Vetencourt et al., 2008; Pizzorusso et al., 2006; Saiepour et al., 2015). During development, immature inhibitory networks shape the CP. In adults, reducing GABAergic inhibition can reopen plasticity windows (Hensch, 2018). Among inhibitory interneurons, parvalbumin-positive (PV) cells are particularly important (Reh et al., 2020). Using chemogenetic approaches in PV-Cre mice, we selectively decreased or increased PV activity in V1 during two hours of IPSI/CONTRA STMD. PV inhibition enhanced OD shifts in adults, mimicking juvenile plasticity: during IPSI STMD, C/I ratios decreased via increased ipsilateral and decreased contralateral VEPs, while during CONTRA STMD, C/I ratios increased due to stronger contralateral and weaker ipsilateral responses. Conversely, PV activation reduced IPSI STMD-induced shifts and abolished CONTRA STMD-induced shifts. Control experiments confirmed that PV modulation alone, without deprivation, did not alter VEPs. These results indicate that STMD plasticity depends at least in part on PV-mediated GABAergic inhibition, and future studies should investigate contributions of other interneuron subtypes, such as SOM- and VIP-positive cells.

These findings have translational relevance. In humans, STMD of the dominant eye produces a perceptual advantage for the deprived eye lasting several hours (Lunghi et al., 2011, 2013), enhanced by physical activity (Lunghi & Sale, 2015). Clinically, brief repeated deprivation of the amblyopic eye combined with moderate exercise improves visual acuity and stereopsis in adult anisometropic amblyopes (Lunghi et al., 2019). Our results support the potential for temporally brief, non-invasive interventions to harness adult cortical plasticity in amblyopia and other contexts.

## Acknowledgements

Alessandro Sale’s laboratory is supported by a *Progetti di ricerca@CNR* grant (grant agreement *Television*). Alessandro Sale also acknowledges funding from the European Union – Next Generation EU, within the framework of the National Recovery and Resilience Plan (NRRP), Investment PE8 – Project *Age-It: “Aging Well in an Aging Society”* (DM 1557, 11.10.2022). This work was co-financed by the European Union – Next Generation EU. We thank Francesca Biondi, Elena Novelli, Renzo Di Renzo, Elena Orsucci and Barbara Roncolini, for their valuable support with animal facility management, technical assistance, and administrative support. We are particularly grateful to Prof. Concetta Morrone for her insightful comments on the manuscript.

## Author contributions

A.S. and N.B. conceived and designed the study. I.D.M. contributed to conceive the experiments, performed all experiments and analyses. G.S. contributed to in vivo electrophysiology and chemogenetics. A.S. and I.D.M. wrote the manuscript with input from all authors. All authors read and approved the final version of the manuscript.

## Supplementary information

### V1 interhemispheric crosstalk modulates OD shifts after short-term monocular deprivation

To investigate the interhemispheric contribution to STMD effects, we blocked V1 activity in the hemisphere contralateral to the one in which electrophysiological recordings were performed in adult C57BL/6J mice, using animals that had previously received intracortical injections of a Cre-dependent viral vector (AAV8-hSyn-HA-hM4D(Gi)-mCherry; **Supplementary Figure 1A,B**). With this procedure, the contralateral V1 was chemogenetically inhibited during the 2 hours of MD.

The animals (n = 5) were recorded on three separate days. On the first day, we assessed the effects of V1 blockade during IPSI STMD (**Supplementary Figure 1A,C**). On the second day, we tested the effects of V1 blockade during CONTRA STMD (**Supplementary Figure 1A,I**). On the third day, all animals underwent three consecutive recordings spaced by 2 hours, during which V1 was blocked for 2 hours but no eye deprivation was applied, to control for any influence of the V1 blockade itself (**Supplementary Figure 1A,O**).

### V1 block and IPSI eye deprivation

Pre-deprivation measurement yielded a C/I VEP ratio of 2.842 ± 0.025. IPSI STMD, coupled with chemogenetic blockade of the contralateral V1, led to a significant reduction in the C/I VEP ratio (One-Way RM ANOVA, DF = 2, F = 256.9; Tukey method, Pre = 2.842 ± 0.025 vs. Post = 1.412 ± 0.086, p < 0.001). Notably, after 2 hours of binocular viewing, the C/I VEP ratio returned to baseline levels (One-Way RM ANOVA, Pre = 2.842 ± 0.025 vs. After 2h = 2.856 ± 0.007, p = 0.978; **Supplementary Figure 1D).**

To dissect the relative contributions of each eye, we analyzed individual VEP amplitudes. STMD induced a significant decrease in CONTRA VEP amplitudes (Two-Way RM ANOVA, eye × time, DF = 2, F = 20.08; Tukey method, Pre = 152.786 ± 13.16 μV vs. Post = 106.748 ± 11.995 μV, p < 0.001; **Supplementary Figure 1F,H**) and a concurrent significant increase in IPSI VEP amplitudes (Pre = 53.976 ± 4.782 μV vs. Post = 77.1 ± 8.164 μV, p < 0.05; **Supplementary Figure 1G,H**). Pre-deprivation, CONTRA responses were significantly stronger than IPSI ones (Contra Pre = 152.786 ± 13.16 μV vs. Ipsi Pre = 53.976 ± 4.782 μV, p < 0.001), consistent with normal OD. Post-deprivation, the responses became closer (Contra Post = 106.748 ± 11.995 μV vs. Ipsi Post = 77.1 ± 8.164 μV, p = 0.0498), reflecting a temporary reduction in CONTRA dominance. After 2 hours of restored binocular vision, VEP amplitudes reverted to their original asymmetry (Contra After 2h = 150.3 ± 14.053 μV vs. Ipsi After 2h = 52.902 ± 5.072 μV, p < 0.001; **Supplementary Figure 1E,H**).

Although STMD still shifted OD toward the deprived eye, the effect was significantly reduced compared to adult wild-type mice subjected to IPSI STMD with an intact contralateral V1. Indeed, the post-STMD C/I VEP ratio was higher in animals with V1 blockade than in wild-types (two-tailed unpaired t-test, Post wild-type = 1.101 ± 0.093 vs. Post V1 block = 1.412 ± 0.086; t = - 3.142, DF = 27, p < 0.05; **Supplementary Figure 2A**).

### V1 block and CONTRA eye deprivation

Baseline OD (C/I VEP ratio) before CONTRA STMD was 2.842 ± 0.066. After STMD, we observed a significant increase in the C/I VEP ratio (One-Way RM ANOVA, DF = 2, F = 186.5; Tukey method, Pre = 2.842 ± 0.066 vs. Post = 3.627 ± 0.064, p < 0.001). This effect was transient, with the ratio returning to baseline after 2 hours of binocular vision (One-Way RM ANOVA, Pre = 2.842 ± 0.066 vs. After 2h = 2.806 ± 0.049, p = 0.976; **Supplementary Figure 1J**).

To determine the source of this shift, we analyzed VEP amplitudes for each eye. Post-deprivation, CONTRA VEP amplitudes increased significantly (Two-Way RM ANOVA, eye × time, DF = 2, F = 17.62; Tukey method, Pre = 126.456 ± 6.925 μV vs. Post = 158.104 ± 13.981 μV, p < 0.01; **Supplementary Figure 1L,N**), whereas IPSI VEP amplitudes did not change (Pre = 44.778 ± 2.889 μV vs. Post = 43.332 ± 3.261 μV, p = 1.00; **Supplementary Figure 1M,N**). The difference between C/I VEP amplitudes remained significant across all three recordings (Two-Way RM ANOVA, p < 0.001 in all comparisons; **Supplementary Figure 1K,N**).

When CONTRA STMD was performed with the contralateral V1 blocked, an OD shift favoring the deprived eye was still observed, but the effect was significantly reduced compared to adult wild-type mice with an intact contralateral V1. Specifically, the post-STMD C/I VEP ratio was lower in animals with V1 blockade than in wild-types (two-tailed unpaired t-test, Post wild-type = 4.080 ± 0.117 vs. Post V1 block = 3.627 ± 0.064; t = 2.972, DF = 29; p < 0.05; **Supplementary Figure 2B**).

### V1 block without eye deprivation

No changes in the C/I VEP ratio were observed across the three recording sessions (One-Way RM ANOVA, Pre = 2.917 ± 0.027, Post = 2.883 ± 0.019, After 2h = 2.862 ± 0.015; DF = 2, F = 1.148; Tukey method, p = 0.364; **Supplementary Figure 1P**). Thus, multiple electrophysiological recordings conducted every 2 hours, during which the contralateral V1 was blocked, did not result in any significant reduction in either CONTRA (Two-Way RM ANOVA, eye × time, DF = 2, F = 0.004; Tukey method, Pre = 103.88 ± 17.612 μV vs. Post = 105.78 ± 18.072 μV, p = 0.999; **Supplementary Figure 1R**) or IPSI VEP amplitudes (Pre = 35.528 ± 6.051 μV vs. Post = 36.682 ± 6.290 μV, p = 1.000; Figure 5R). The difference between C/I VEP amplitudes remained significant across all three recordings (Two-Way RM ANOVA, p < 0.01 in all comparisons; **Supplementary Figure 1Q,R**).

These results reveal that V1 interhemispheric crosstalk partially drives STMD-induced homeostatic plasticity, as blocking the non-recorded V1 significantly attenuates effects compared to wild-type mice. Yet, this rapid form of homeostatic plasticity persists even with functional silencing of the contralateral hemisphere.

## Supplementary figure legends

**Supplementary Figure 1 Impact of V1 interhemispheric crosstalk on IPSI/CONTRA short-term monocular deprivation (STMD).**

**a,** Schematic illustration of the experimental protocol. In the upper left, a coronal section of the mouse brain is shown. The boundaries of V1 in both hemispheres are outlined in black. In the right hemisphere’s V1, the green area indicates the monocular zone, while the yellow area represents the binocular zone, where electrophysiological recordings were conducted. The viral injection was administered in the left V1, highlighted in red, which corresponds to the hemisphere contralateral to the recording site. **b,** Coronal section of the mouse brain zoomed in on V1, with the border outlined by a white dashed line, illustrating that viral expression was confined within the V1 boundaries. In the upper right corner, brain coordinates are indicated: dorsal (D) to ventral (V) and anterior (A) to posterior (P). The red line marks the location of the coronal section shown. **c,** Timeline of electrophysiological recordings on Day 1: baseline recordings before IPSI STMD (Pre), recordings immediately after STMD (Post), and recordings following two hours of restored binocular vision (After 2h). **d,** Contralateral/Ipsilateral (C/I) VEP ratio in response to a low spatial frequency grating stimulus (0.06 c/deg). The C/I VEP ratio significantly decreases immediately after IPSI STMD compared to baseline (One-way ANOVA RM, p<0.001), and returns to baseline levels following two hours of binocular vision (One-way RM ANOVA, p=0.978). **e,** Mean VEP amplitudes recorded from contralateral and ipsilateral eyes at all timepoints. In the upper-right, mean VEP amplitudes from the contralateral eye before and after IPSI STMD. Two hours of monocular deprivation results in a significant reduction in contralateral VEP amplitudes (Two-way ANOVA RM, p<0.001). The darkest line represents the mean. In the bottom-right, mean VEP amplitudes from the ipsilateral eye before and after IPSI STMD. Two hours of monocular deprivation induces a significant increase in ipsilateral VEP amplitudes (Two-way ANOVA RM, p<0.05). The darkest line represents the mean. **f,** Representative VEP traces from one mouse, demonstrating changes across Pre and Post STMD. **g,** Timeline of electrophysiological recordings on Day 2: baseline recordings before CONTRA STMD (Pre), recordings immediately after STMD (Post), and recordings following two hours of restored binocular vision (After 2h). **h,** The C/I VEP ratio significantly increases immediately after CONTRA STMD compared to baseline (One-way ANOVA RM, p<0.001), and returns to baseline levels following two hours of binocular vision (One-way RM ANOVA, p=0.978). **i,** Mean VEP amplitudes recorded from contralateral and ipsilateral eyes at all timepoints. In the upper-right, mean VEP amplitudes from the contralateral eye before and after CONTRA STMD. Two hours of monocular deprivation results in a significant increase in contralateral VEP amplitudes (Two-way ANOVA RM, p<0.01). The darkest line represents the mean. In the bottom-right, mean VEP amplitudes from the ipsilateral eye before and after CONTRA STMD. Two hours of monocular deprivation induces a non-significant decrease in ipsilateral VEP amplitudes (Two-way ANOVA RM, p=1.00). The darkest line represents the mean. **j,** Representative VEP traces from one mouse, demonstrating changes across Pre and Post STMD. **k,** Timeline of electrophysiological recordings on Day 3. **l,** Three consecutive electrophysiological recordings without monocular deprivation did not produce significant changes in the C/I VEP ratio across timepoints (One-way ANOVA RM, p=0.364). **m,** Mean VEP amplitudes recorded from contralateral and ipsilateral eyes at all timepoints. **n,** Representative VEP traces from one mouse at baseline (Pre) and after two hours of binocular vision (Post).

C: contralateral eye; I: ipsilateral eye; Pre: before STMD; Post: after STMD; After: two hours after the end of monocular deprivation.

Statistical significance: ns: not significant; *p < 0.05; **p < 0.01; **p < 0.001; error bars indicate S.E.M.

**Supplementary Figure 2 Comparative evaluation of IPSI/CONTRA short-term monocular deprivation (STMD) effects in adult naïve mice and those with inactivation of the non-recorded V1.**

**a,** Inactivation of V1 in the non-recorded hemisphere during the deprivation phase significantly reduced the effect of IPSI STMD compared to naïve mice (two-tailed unpaired t-test, Post naïve vs. Post block V1; p<0.05). **b,** Following CONTRA STMD, mice with V1 inactivation in the non-recorded hemisphere during STMD showed a reduced effect (two-tailed unpaired t-test, Post naïve vs. Post block V1; p<0.05).

Group definitions: Naïve = C57BL/6J adult mice; Block V1 = mice with suppressed activity in the non-recorded V1 during deprivation.

Statistical significance: *p < 0.05; **p < 0.01; **p < 0.001; error bars indicate S.E.M.

## References

1. Antonini, A., Stryker, M. P., & Keck, W. M. (1998). Effect of sensory disuse on geniculate afferents to cat visual cortex.

2. Barnes, S. J., Sammons, R. P., Jacobsen, R. I., Mackie, J., Keller, G. B., & Keck, T. (2015). Subnetwork-Specific Homeostatic Plasticity in Mouse Visual Cortex In Vivo. Neuron, 86(5), 1290–1303. 10.1016/j.neuron.2015.05.010

3. Binda, P., & Lunghi, C. (2017). Short-Term Monocular Deprivation Enhances Physiological Pupillary Oscillations. Neural Plasticity, 2017. 10.1155/2017/6724631

4. Chen, L., Li, X., Tjia, M., & Thapliyal, S. (2022). Homeostatic plasticity and excitation-inhibition balance: The good, the bad, and the ugly. In Current Opinion in Neurobiology (Vol. 75). Elsevier Ltd. 10.1016/j.conb.2022.102553

5. Espinosa, J. S., & Stryker, M. P. (2012). Development and Plasticity of the Primary Visual Cortex. In Neuron (Vol. 75, Issue 2, pp. 230–249). 10.1016/j.neuron.2012.06.009

6. Fagiolini, M., Fritschy, J.-M., Löw, K., Möhler, H., Rudolph, U., & Hensch, T. (2004). Specific GABAA Circuits for Visual Cortical Plasticity. Science, 303, 1681–1683.

7. Fagiolini, M., & Hensch, T. K. (2000). Inhibitory threshold for critical-period activation in primary visual cortex. Nature, 404, 183–186.

8. Gordon, J. A., Sttyker, M. P., Program, N. G., & Keck, W. M. (1996). Experience-Dependent Plasticity of Binocular Responses in the Primary Visual Cortex of the Mouse. In The Journal of Neuroscience (Vol. 76, Issue 10).

9. Hensch, T. K. (2003). Controlling the critical period. Neuroscience Research, 47(1), 17–22. 10.1016/S0168-0102(03)00164-0

10. Hensch, T. K. (2005). Critical period plasticity in local cortical circuits. In Nature Reviews Neuroscience (Vol. 6, Issue 11, pp. 877–888). 10.1038/nrn1787

11. Hensch, T. K. (2018). Critical Periods in Cortical Development. In The Neurobiology of Brain and Behavioral Development (pp. 133–151). Elsevier Inc. 10.1016/B978-0-12-804036-2.00006-6

12. Hensch, T. K., Fagiolini, M., Mataga, N., Stryker, M. P., Baekkeskov, S., & Kash, S. F. (1998). Local GABA Circuit Control of Experience-Dependent Plasticity in Developing Visual Cortex. In Proc. Natl. Acad. Sci. U.S.A (Vol. 137). Patel. http://science.sciencemag.org/

13. Hofer, S. B., Mrsic-Flogel, T. D., Bonhoeffer, T., & Hübener, M. (2006). Lifelong learning: ocular dominance plasticity in mouse visual cortex. In Current Opinion in Neurobiology (Vol. 16, Issue 4, pp. 451–459). 10.1016/j.conb.2006.06.007

14. Hubel, D. H., & Wiesel, T. N. (1959). Receptive fields of single neurones in the cat’s striate cortex. The Journal of Physiology, 148(3), 574. 10.1113/JPHYSIOL.1959.SP006308

15. Hubel, D. H., & Wiesel, T. N. (1970). THE PERIOD OF SUSCEPTIBILITY TO THE PHYSIOLOGICAL EFFECTS OF UNILATERAL EYE CLOSURE IN KITTENS. In J. Physiol (Vol. 206).

16. Kaneko, M., Stellwagen, D., Malenka, R. C., & Stryker, M. P. (2008). Tumor Necrosis Factor-α Mediates One Component of Competitive, Experience-Dependent Plasticity in Developing Visual Cortex. Neuron, 58(5), 673–680. 10.1016/j.neuron.2008.04.023

17. Kaplan, E. S., Cooke, S. F., Komorowski, R. W., Chubykin, A. A., Thomazeau, A., Khibnik, L. A., Gavornik, J. P., & Bear, M. F. (2016). Contrasting roles for parvalbumin-expressing inhibitory neurons in two forms of adult visual cortical plasticity. ELife. 10.7554/eLife.11450.001

18. Kasamatsu, T., & Imamura, K. (2020). Ocular dominance plasticity: Molecular mechanisms revisited. In Journal of Comparative Neurology (Vol. 528, Issue 17, pp. 3039–3074). Wiley-Liss Inc. 10.1002/cne.25001

19. Kawasaki, A., Wisniewski, S., Healey, B., Pattyn, N., Kunz, D., Basner, M., & Münch, M. (2018). Impact of long-term daylight deprivation on retinal light sensitivity, circadian rhythms and sleep during the Antarctic winter. Scientific Reports, 8(1). 10.1038/s41598-018-33450-7

20. Lehmann, K., & Löwel, S. (2008). Age-dependent ocular dominance plasticity in adult mice. PLoS ONE, 3(9). 10.1371/journal.pone.0003120

21. Levelt, C. N., & Lubener, M. (2012). Critical-period plasticity in the visual cortex. In Annual Review of Neuroscience (Vol. 35, pp. 309–330). 10.1146/annurev-neuro-061010-113813

22. Lunghi, C., Berchicci, M., Morrone, M. C., & Di Russo, F. (2015). Short-term monocular deprivation alters early components of visual evoked potentials. Journal of Physiology, 593(19), 4361–4372. 10.1113/JP270950

23. Lunghi, C., Burr, D. C., & Concetta Morrone, M. (2013). Long-term effects of monocular deprivation revealed with binocular rivalry gratings modulated in luminance and in color. Journal of Vision, 13(6). 10.1167/13.6.1

24. Lunghi, C., Burr, D. C., & Morrone, C. (2011). Brief periods of monocular deprivation disrupt ocular balance in human adult visual cortex. In Current Biology (Vol. 21, Issue 14). 10.1016/j.cub.2011.06.004

25. Lunghi, C., Emir, U. E., Morrone, M. C., & Bridge, H. (2015). Short-Term monocular deprivation alters GABA in the adult human visual cortex. Current Biology, 25(11), 1496–1501. 10.1016/j.cub.2015.04.021

26. Lunghi, C., & Sale, A. (2015). A cycling lane for brain rewiring. In Current Biology (Vol. 25, Issue 23, pp. R1122–R1123). Cell Press. 10.1016/j.cub.2015.10.026

27. Lunghi, C., Sframeli, A. T., Lepri, A., Lepri, M., Lisi, D., Sale, A., & Morrone, M. C. (2019). A new counterintuitive training for adult amblyopia. Annals of Clinical and Translational Neurology, 6(2), 274–284. 10.1002/acn3.698

28. Maya Vetencourt, J. F., Sale, A., Viegi, A., Baroncelli, L., De Pasquale, R., O’Leary, O. F., Castrén, E., & Maffei, L. (2008). The Antidepressant FluoxetineRestores Plasticity in the Adult Visual Cortex. Science, 320(5874), 385–388. 10.1126/science.1155307

29. Mazziotti, R., Baroncelli, L., Ceglia, N., Chelini, G., Sala, G. Della, Magnan, C., Napoli, D., Putignano, E., Silingardi, D., Tola, J., Tognini, P., Arthur, J. S. C., Baldi, P., & Pizzorusso, T. (2017). Mir-132/212 is required for maturation of binocular matching of orientation preference and depth perception. Nature Communications, 8. 10.1038/ncomms15488

30. Menicucci, D., Lunghi, C., Zaccaro, A., Morrone, M. C., & Gemignani, A. (2022). Mutual interaction between visual homeostatic plasticity and sleep in adult humans. ELife, 11. 10.7554/eLife.70633

31. Mrsic-Flogel, T. D., Hofer, S. B., Ohki, K., Reid, R. C., Bonhoeffer, T., & Hübener, M. (2007). Homeostatic Regulation of Eye-Specific Responses in Visual Cortex during Ocular Dominance Plasticity. Neuron, 54(6), 961–972. 10.1016/j.neuron.2007.05.028

32. Nguyen, B. N., Srinivasan, R., & McKendrick, A. M. (2023). Short-term homeostatic visual neuroplasticity in adolescents after two hours of monocular deprivation. IBRO Neuroscience Reports, 14, 419–427. 10.1016/j.ibneur.2023.04.003

33. Petros, T. J., Rebsam, A., & Mason, C. A. (2008). Retinal axon growth at the optic chiasm: To cross or not to cross. In Annual Review of Neuroscience (Vol. 31, pp. 295–315). 10.1146/annurev.neuro.31.060407.125609

34. Pietrasanta, M., Restani, L., & Caleo, M. (2012). The Corpus Callosum and the Visual Cortex: Plasticity Is a Game for Two. Neural Plasticity, 2012(1), 838672. 10.1155/2012/838672

35. Pizzorusso, T., Medini, P., Landi, S., Baldini, S., Berardi, N., & Maffei, L. (2006). Structural and functional recovery from early monocular deprivation in adult rats. www.pnas.orgcgi10.1073pnas.0602657103

36. Porciatti, V., Pizzorusso, T., & Maffei, L. (1999). The visual physiology of the wild type mouse determined with pattern VEPs. In Vision Research (Vol. 39). www.elsevier.com

37. Prosper, A., Pasqualetti, M., Morrone, M. C., & Lunghi, C. (2023). The duration effect of short-term monocular deprivation measured by binocular rivalry and binocular combination. Vision Research, 211. 10.1016/j.visres.2023.108278

38. Ranson, A., Cheetham, C. E. J., Fox, K., & Sengpiel, F. (2012). Homeostatic plasticity mechanisms are required for juvenile, but not adult, ocular dominance plasticity. Proceedings of the National Academy of Sciences of the United States of America, 109(4), 1311–1316. 10.1073/pnas.1112204109

39. Reh, R. K., Dias, B. G., Nelson, C. A., Kaufer, D., Werker, J. F., Kolbh, B., Levine, J. D., & Hensch, T. K. (2020). Critical period regulation acrossmultiple timescales. In Proceedings of the National Academy of Sciences of the United States of America (Vol. 117, Issue 38, pp. 23242–23251). National Academy of Sciences. 10.1073/pnas.1820836117

40. Restani, L., Cerri, C., Pietrasanta, M., Gianfranceschi, L., Maffei, L., & Caleo, M. (2009). Functional Masking of Deprived Eye Responses by Callosal Input during Ocular Dominance Plasticity. Neuron, 64(5), 707–718. 10.1016/j.neuron.2009.10.019

41. Rittenhouse, C. D., Shouval, H. Z., Paradiso, M. A., & Bear, M. F. (1999). Monocular deprivation induces homosynaptic long-term depression in visual cortex. Nature, 347–350.

42. Saiepour, M. H., Rajendran, R., Omrani, A., Ma, W. P., Tao, H. W., Heimel, J. A., & Levelt, C. N. (2015). Ocular dominance plasticity disrupts binocular inhibition-excitation matching in visual cortex. Current Biology, 25(6), 713–721. 10.1016/j.cub.2015.01.024

43. Sato, M., & Stryker, M. P. (2008). Distinctive features of adult ocular dominance plasticity. Journal of Neuroscience, 28(41), 10278–10286. 10.1523/JNEUROSCI.2451-08.2008

44. Sawtell, N. B., Frenkel, M. Y., Philpot, B. D., Nakazawa, K., Tonegawa, S., & Bear, M. F. (2003). NMDA Receptor-Dependent Ocular Dominance Plasticity in Adult Visual Cortex of primary visual cortex in anesthetized animals. VEPs reflect population synaptic currents, and the ratio of VEP amplitudes evoked by alternating patterned visual. In Neuron (Vol. 38).

45. Steinwurzel, C., Morrone, M. C., Sandini, G., & Binda, P. (2023). Active vision gates ocular dominance plasticity in human adults. Current Biology, 33(20), R1038–R1040. 10.1016/j.cub.2023.08.062

46. Tong, F., Meng, M., & Blake, R. (2006). Neural bases of binocular rivalry. In Trends in Cognitive Sciences (Vol. 10, Issue 11, pp. 502–511). 10.1016/j.tics.2006.09.003

47. Turrigiano, G. G. (2017). The dialectic of hebb and homeostasis. Philosophical Transactions of the Royal Society B: Biological Sciences, 372(1715). 10.1098/rstb.2016.0258

48. Turrigiano, G. G., & Nelson, S. B. (2004). Homeostatic plasticity in the developing nervous system. In Nature Reviews Neuroscience (Vol. 5, Issue 2, pp. 97–107). European Association for Cardio-Thoracic Surgery. 10.1038/nrn1327

49. Turrigiano, G., Leslie, K., Desai, N., Rutherford, L., & Nelson, S. (1998). Activity-dependent scaling of quantal amplitude in neocortical neurons.

