## Supplementary figures and images for "Short-term monocular deprivation engages rapid, inhibition-gated ocular dominance plasticity in mouse visual cortex"

### Supplementary Fig. 1

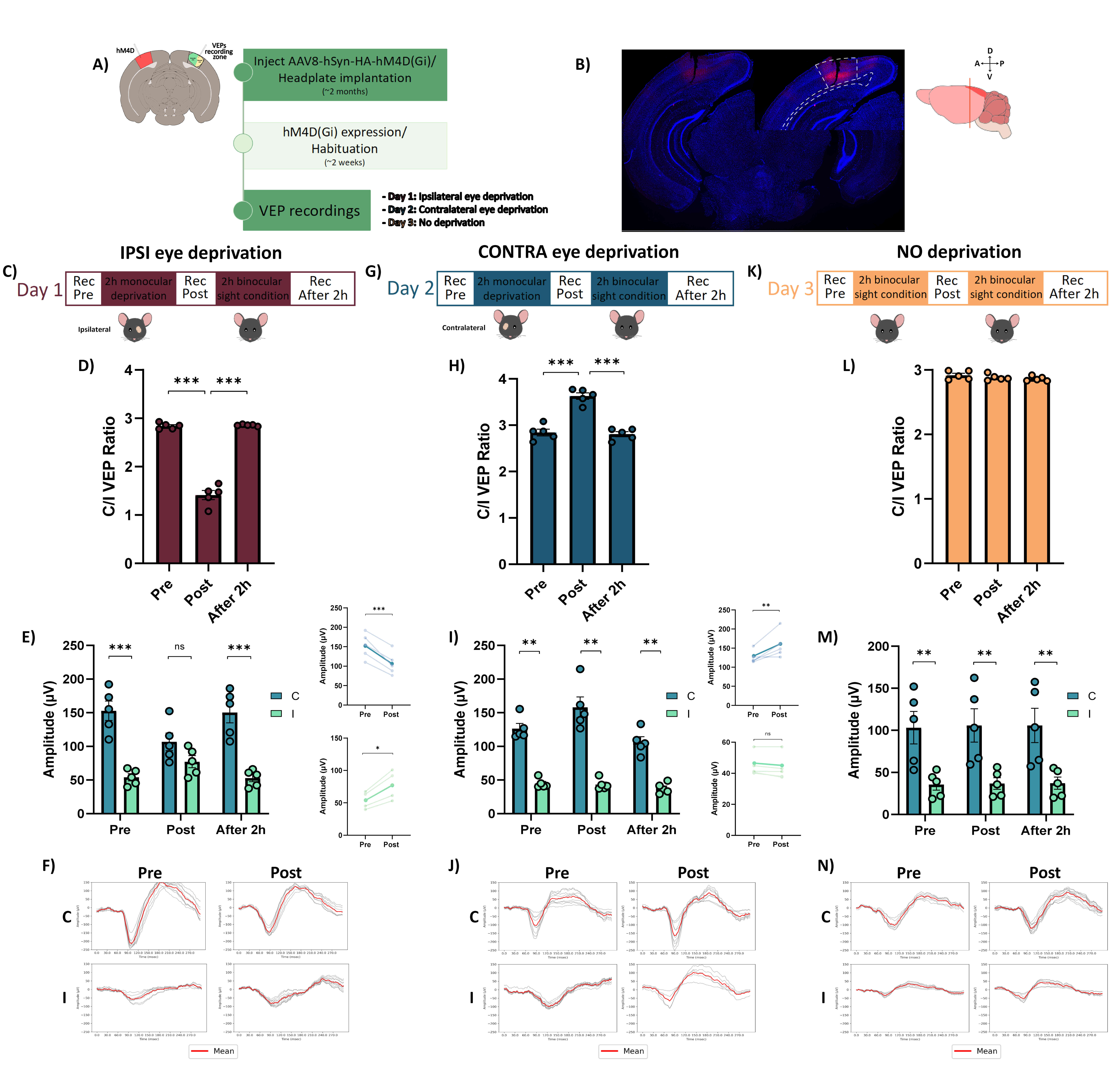

### Supplementary Fig. 2

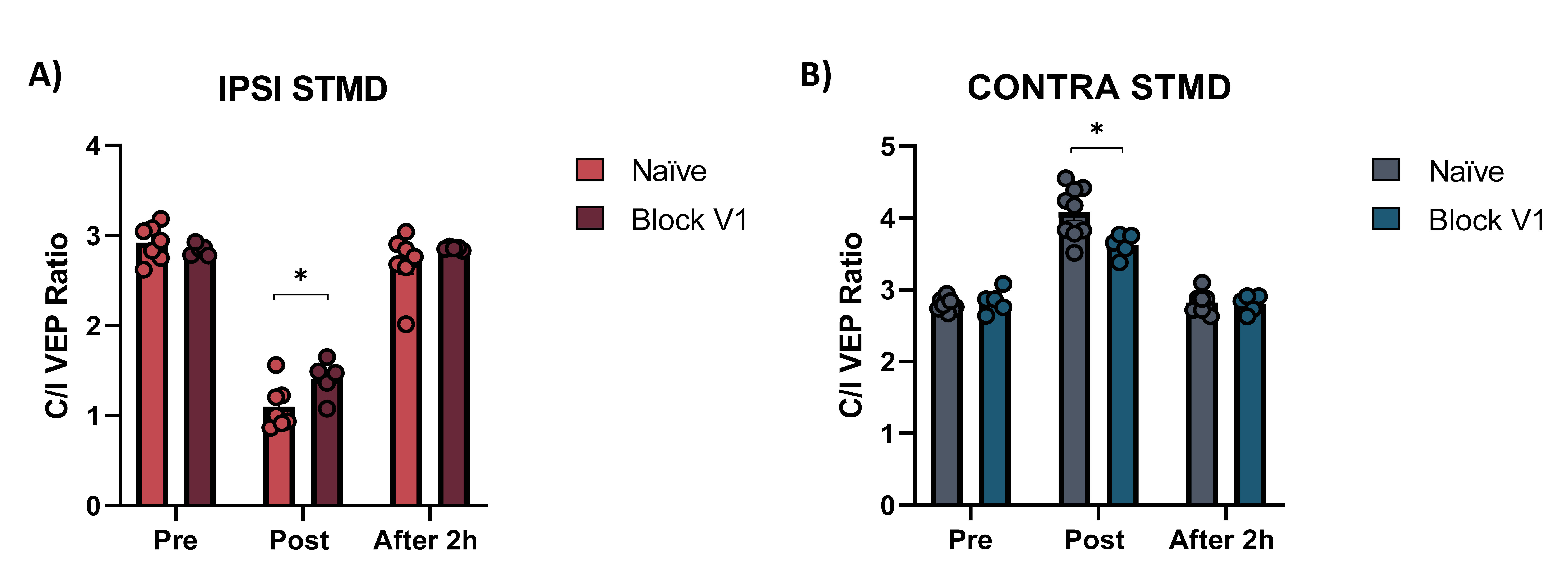
